# Relating sex differences in cortical and hippocampal microstructure to sex hormones

**DOI:** 10.1101/2023.11.01.565213

**Authors:** Svenja Küchenhoff, Şeyma Bayrak, Rachel G. Zsido, Amin Saberi, Boris Bernhardt, Susanne Weis, H. Lina Schaare, Julia Sacher, Simon Eickhoff, Sofie L. Valk

## Abstract

Sex hormone receptors are expressed widely in both neurons and glial cells, which allows them to interact with the brain’s major cell groups via several molecular mechanisms. These mechanisms lead to sex differences in brain structure as well as hormone-induced plasticity in the female brain across the menstrual cycle. Adding to the literature on volumetric changes in cortical structure, here we set out to investigate sex differences in the microstructure of the human cortex in relation to sex hormones. We assessed regional variation in cortical microstructure as a function of sex, hormonal status and sex hormone receptor gene expression distribution based on quantitative intracortical profiling *in vivo* using the magnetic resonance imaging based T1w/T2w ratio in 992 healthy females and males of the Human Connectome Project young adult sample. We demonstrate that microstructure in isocortex and hippocampus differs regionally between males and females, that this effect varies with hormone levels of females and that implicated brain regions overlap with estrogen receptor and sex steroid synthesis gene expression. Lastly, we show that sex and sex hormone related brain structure variations are most pronounced in areas of low laminar cortical complexity (agranular cortex), which are also predicted to be most plastic based on their cytoarchitectural properties. Together, our data thus are suggestive of sex differences in cortical and hippocampal microstructure, as well as the modulatory function of sex hormones on these measures. Albeit correlative, this study underscores the importance of incorporating sex hormone variables into the investigation of brain structure and plasticity.

## Introduction

Brain structure constantly adapts to both environmental and bodily demands. One regulatory mechanism allowing for body-brain interactions via structural plasticity are circulating sex hormones produced in the gonads, which the brain controls via the pituitary gland (McEwen, 2012). This type of endocrine structural plasticity has been repeatedly demonstrated on fast (hours-to-days) and slow (years) timescales, with animal research providing robust evidence on a molecular basis (Barha & Galea, 2010; Been et al., 2022; Cooke & Woolley, 2005; Hara et al., 2015; Patel et al., 2013; Woolley & McEwen, 1993). Pioneering work in a more challenging human experimental setting has presented first evidence for endocrinologically linked plasticity effects in the human brain. Building on these macro-level findings presenting volumetric changes in the human cortex (Seitz et al., 2019; Taylor et al., 2020; Witte et al., 2010; Zsido et al., 2023), here we investigate the underlying intracortical microstructural changes of sex and sex hormones. In an effort to bridge traditional neuroanatomy and neuroimaging, we built several intracortical measures *in vivo* based on the ratio of T1- over T2 weighted (T1w/T2w) MRI intensities, and investigated if intra-individual differences in these proxies could be linked to male-female sex differences. Since biological sex is a multidimensional construct (*Measuring Sex, Gender Identity, and Sexual Orientation | The National Academies Press*, n.d.), we furthermore investigate the role of sex hormones as one dimension of sex in these structural differences.

Sex hormone receptors are expressed in neurons as well as glial cells, which allows sex hormones to influence myelination (Schumacher et al., 2012), dendritic spine morphology and density (de Castilhos et al., 2008; Romeo et al., 2004), and cell metabolism (Brinton, 2009), via ion channels, second-messenger systems and gene expression in the brain (McEwen, 2012). The pattern of sex hormone synthesis products as well as sex hormone receptors in the brain differs by region, changes over the lifetime, and diverges between self reported males and females^1^ (Moraga-Amaro et al., 2018; Quadros et al., 2002; Sharma & Thakur, 2006; Wilson et al., 2011). The influence of sex hormones on brain structure is strongest during critical phases in development such as early perinatal development and puberty, where the expression of sex hormones trigger the emergence of sexually divergent traits (Arnold & Breedlove, 1985). Sex hormones, however, continue to modulate brain structure throughout adulthood (Rehbein et al., 2021; Romeo et al., 2004). In ovulating individuals, brain structure varies on a short time-scale of days to weeks alongside sex steroid fluctuations during the menstrual cycle(Barth et al., 2016; Fernández et al., 2003; Protopopescu et al., 2008; Zsido et al., 2023), where the progesterone levels increase up to 80-fold, and estradiol levels up to 8-fold over a period of 25-34 days (Mihm et al., 2011; Stricker et al., 2006). Additionally, brain structure can also be modulated by exogenous sex hormones which influence sex-hormone profiles such as the commonly prescribed oral contraceptive pill (OC) in the medium term of weeks to months (Petersen et al., 2023).

Overall, the links between sex hormones and volumetric changes in brain structure on a short, medium and long timescale are well documented. For example, macroscale structural changes on the short time scale (weeks) covary with the menstrual cycle, where paralimbic brain structures in particular adjust their structure to fluctuations of estrogen and progesterone (De Bondt et al., 2013; Dubol et al., 2021; Lisofsky et al., 2015; Ossewaarde et al., 2013; Pletzer et al., 2018). On a the scale of months, the use of OC in comparison to naturally cycling females has been shown to decrease gray matter of the amygdala and the parahippocampal gyrus (Lisofsky et al., 2016), and the cortical thickness of the prefrontal cortex (Petersen et al., 2021, 2023). Furthermore, the intense hormone level changes during pregnancy go along with volumetric changes in medial temporal and medial prefrontal areas relevant for social cognition (Hoekzema et al., 2017). Sex differences in gray matter volume in adults that developed over years from the onset of diverging sex hormonal profiles in puberty, are partly explained by circulating testosterone, progesterone and 17β-estradiol levels (Lentini et al., 2013; Witte et al., 2010). While all of these results refer to volumetric changes measured with structural magnetic resonance imaging (MRI), human microstructural sex differences *in-vivo* have not yet been characterized, and it remains elusive if sex hormones might play a role in these variations. Contradicting reports on volumetric cortical sex differences have further been subject to critiques about insufficient control of systematic differences in brain size (DeCasien et al., 2022). An analysis of sex differences on the microstructural level will yield a more nuanced characterization of sex differences free from these biases.

Multiple neuroanatomical accounts have illustrated the intrinsic link between microstructural properties, inherent brain organization principles, and brain function (Barbas, 1986, 1997; Douglas & Martin, 2004; Langdon et al., 2023). Microstructural changes within the cortical sheath are traditionally examined *post mortem* using cell-staining procedures (*Brodmann: Die Rindenfelder Der Niederen Affen - Google Scholar*, n.d.; von Economo & Koskinas, 1925; Zilles et al., 2002). On this micro-level, the human cortex is structured into several cell layers. The amount and prominence of each layer as well as the sharpness of their boundaries varies across the cortex, so that cortical areas can be classified into different types according to their laminar elaboration (García-Cabezas et al., 2019, 2020; von Economo & Koskinas, 1925). These variations in cortical types are systematically linked to the cortex’ inherent property of plasticity (García-Cabezas et al., 2017, 2019), such that simpler laminar structures (e.g. paralimbic structures) are hypothesized to be more plastic than highly elaborate areas (e.g. primary visual cortex) (García-Cabezas et al., 2017; Sasaki et al., 2015). Amongst others, one explanatory factor for this covariation of laminar differentiation with plasticity is the amount of intracortical myelin, which inhibits plasticity in the brain (Akbik et al., 2012; Boghdadi et al., 2018; Glasser et al., 2014; McGee et al., 2005; Monje, 2018; Raiker et al., 2010). Intracortical myelin content correlates with laminar differentiation so that more elaborate laminar architecture is characterized by higher intracortical myelin content and higher stability (García-Cabezas et al., 2017; Glasser & Van Essen, 2011). Lastly, gradients of microstructural variation running along major axes of organization in the cortex support variation in brain function (Kharabian Masouleh et al., 2020; Vezoli et al., 2021; Waymel et al., 2020). Thus, examining variations in i) microstructural tissue properties, ii) cortical lamination and iii) the microstructural inter-regional organization *in-vivo* in relation to endocrine plasticity may yield evidence about the basis of macroscale plasticity observed with neuroimaging.

To target this question, in the present study, we thus investigated sex differences in microstructural variations and characterized to what extent sex hormones might be linked to the identified effects. We studied cortical microstructure with quantitative profiling of intracortical properties based on the MRI T1w/T2w ratio. More precisely, we analyzed i) an average measure of regional intracortical tissue properties, ii) a measure of the local weighting of upper vs. lower cortical layers, and iii) a measure of the relative distribution of microstructural organization across the cortex. We leveraged n = 1093 T1w/T2w MRI scans from the HCP young adult dataset and quantified these three local and global properties of individual intracortical microstructure across the cortex.

We then contrasted these microstructural measures between females and males, tested how these sex-differences vary if systematically comparing males with females of particular hormonal profiles (approximated by self-reported menstrual cycle phase and OC use) and quantified how these effects overlap with transcriptomic maps of sex-hormone related genes. Lastly, we linked the observed neuroendocrine plasticity effects to a model from traditional cytoarchitectural neuroscience, which predicts elevated plasticity for areas characterized by less elaborate laminar differentiation.

## Results

### Characterization of the three intracortical T1wT2w profile measures across the whole sample (Fig. 1)

We analyzed the microstructural data of n = 1093 subjects (n = 594 females^2^) from the HCP1200 young adult dataset (Van Essen et al., 2013), that was projected onto the cortical surface and parcellated into 400 Schaefer parcels (**Figure 1**A) (Schaefer et al., 2018). We built three different local and global measures of intracortical microstructure, which focus on different quantitative aspects of the microstructural properties: *i)* the microstructural profile mean, representing the local mean T1w/T2w signal intensity across the cortical gray matter tissue (**Figure 1** D.i), *ii)* the microstructural profile skewness indicating the local dominance of T1w/T2w intensity in superficial layers compared to deeper layers (**Figure 1** D.ii), and *iii)* the principal microstructural gradient, which reflects an organizational axis of T1w/T2w intensity covariation along the cortex (**Figure 1** D.iii). To gain insight on the endocrinological effects of the hippocampal microstructure, we further projected T1w/T2w intensities on an *unfolded* hippocampal formation that was automatically delineated using the HippUnfold toolbox (DeKraker et al., 2018, 2023) (**Figure** 1 C). We included the hippocampal data of n = 867 individuals, for whom data in sufficient quality was available. All measures had previously been validated with histological work (DeKraker et al., 2020; Paquola, Wael, et al., 2019; Zilles et al., 2002). The group-averaged maps of the three cortical microstructure measures (Figure 1 C) and the T1w/T2w signal intensity pattern of hippocampus (Figure 1 D) derived from the present sample broadly overlap with previous microstructural mappings of the human cortex (DeKraker et al., 2018; García-Cabezas et al., 2020; Glasser & Van Essen, 2011; Paquola, Wael, et al., 2019; Paquola et al., 2020; Vos de Wael et al., 2018).

**Figure 1.**
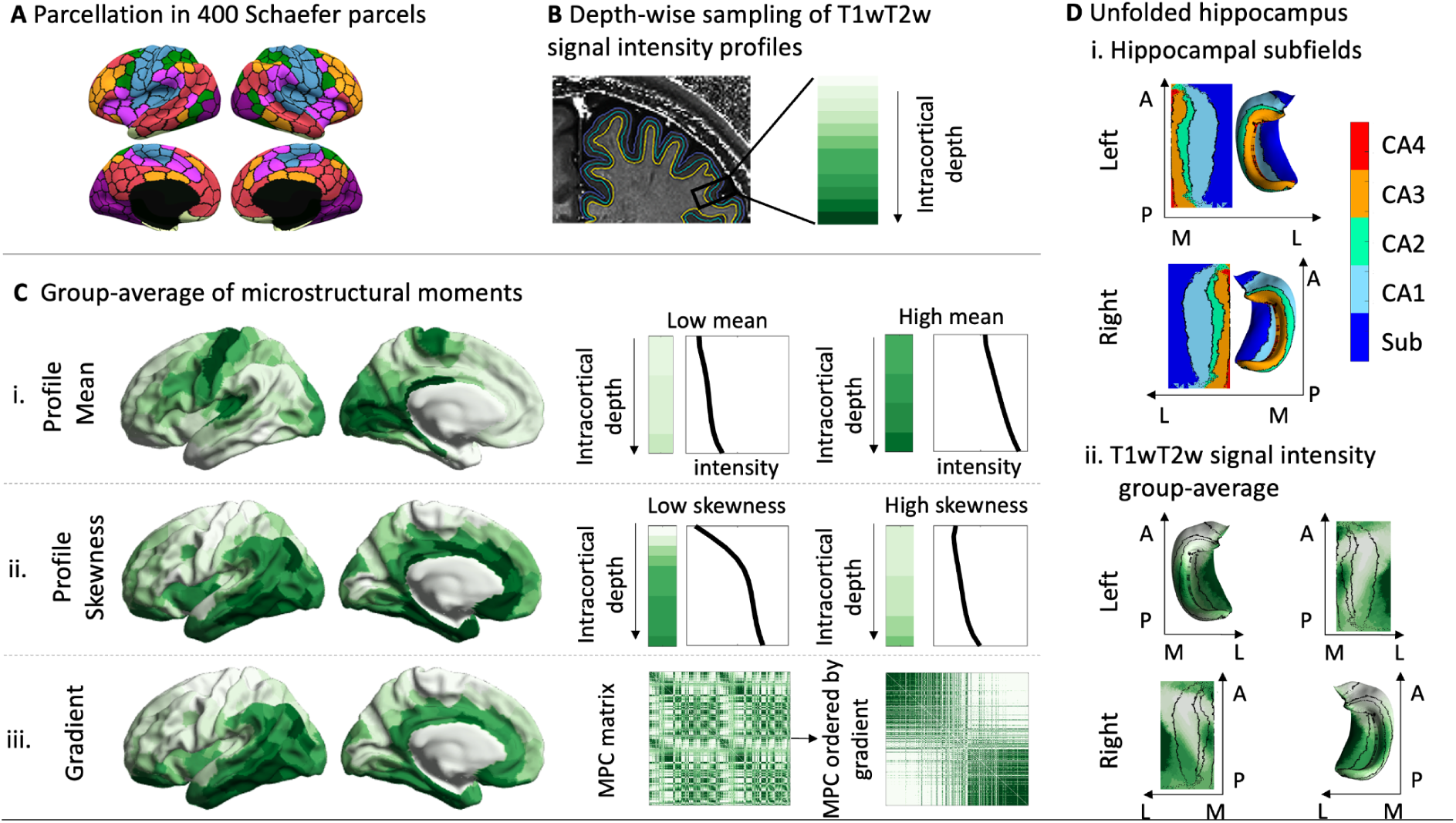
Intracortical T1w/T2w signal intensity profiling. (A) Parcellation scheme. (B) Intracortical sampling to build microstructural profiles. Twelve equivolumetric surfaces are put between cortical surface and white matter boundary of a single subject, yielding 12 sample points at different intracortical depths. (C) left: Group average of microstructural measures, plotted on the cortical surface. right: i. and ii., examples for parcels with a high and low profile mean and skewness per intracortical sample point, respectively. iii., Microstructural profile covariance matrix (MPC) based on correlations of microstructural profiles between pairs of parcels. In the right MPC, parcels are ordered according to their microstructural differentiation, using the principal component derived from diffusion embedding. (D) i. Map of the hippocampal subfields after extraction and unfolding of the hippocampus, and ii. group-average T1w/T2w signal intensity for the left and right hippocampus. Abbreviations: MPC = microstructural profile covariance matrix, CA = Cornu ammonis, Sub = Subiculum, M = medial, L = lateral, P = posterior, A = anterior

### Intracortical microstructural organization differs between males and females (Fig. 2)

To extract sex from our dataset, we categorized everyone as ‘female’ who self-reported their gender as female and indicated they are or have been menstruating in their lives. To identify differences between males and females in each of our three intracortical microstructure measures, we first modeled each measure as a function of intracranial volume, age and sex, and then computed FDR-corrected two-tailed t-tests for our contrast of interest (females > males) for each parcel separately (*q <* 0.05). To be able to compare effects between microstructural measures, we then transformed t-values to Cohen’s *d* effect size values, where a positive Cohen’s *d* represents parcels that had significantly higher microstructural measure values in women, and negative Cohen’s d represents significantly higher values in males in the respective measure.

Comparing the microstructural profile *mean* between males and females, we found that males on average had a higher T1w/T2w profile mean across the whole cortex (cortex-wide average *mean* _*men*_ = 1.7929, *SD* _*men*_ = 0.1191; cortex-wide average *mean* _*females*_ = 1.7498; *SD*_*females*_ = 0.0979). These differences were particularly pronounced bilaterally in parietal, primary sensory motor areas, and unilaterally in left superior temporal and frontal areas (mean effect size of parcels that were significantly higher for males after FDR correction: *d* = -.3214, *SD*_*all neg parcels*_ = 0.1043, **Figure 2**A). The T1w/T2w profile mean of the entorhinal cortex was slightly higher in females, but this difference did not survive FDR correction. As a region bordering the entorhinal cortex, this pattern extended to the subiculum and the CA1 in the hippocampus, which also showed a higher T1w/T2w profile mean for females (mean positive effect size of all FDR-corrected areas in the hippocampus *d* = .2559, *SD*_*all pos parcels*_ = 0.0397, **Figure 2**B). In the most medial part of the hippocampus this pattern reversed such that the hippocampal subfields CA2 and CA3 had a higher profile mean in males than in females (mean negative effect size of all FDR-corrected areas *d* = −0.3315, *SD*_*all neg parcels*_ = 0.1054). The effects were slightly more pronounced in the right than in the left hippocampus.

**Figure 2.**
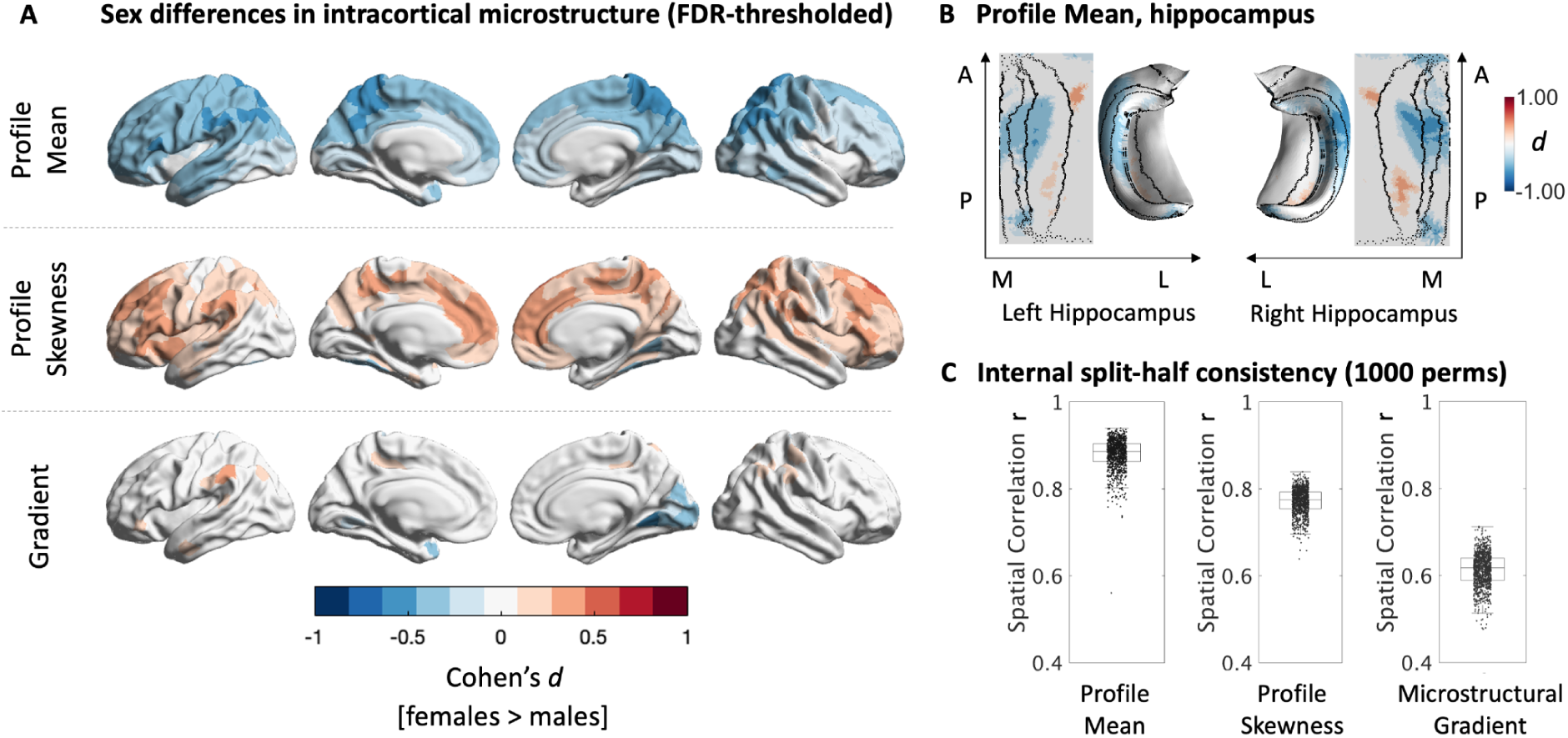
Sex differences in the brain’s microstructural organization. (A) FDR-thresholded Cohen’s *d* maps showing significant sex differences (females-males) in intracortical microstructure: T1w/T2w based intracortical profile mean, profile skewness and the microstructural gradient. Red colors represent microstructural values were higher for females, blue represent values higher for men. (B) 2D and 3D FDR thresholded effect-size maps of the unfolded hippocampus, showing significant differences in T1w/T2s mean between females and men. (C) Boxplots represent Pearson’s r-values between unthresholded t-statistics resulting from two respective split-halves of the sample (n = 1000 permutations) comparing profile mean, profile skewness and microstructural gradient between females and males, indicating the reliability of each measure.

Sex differences in T1w/T2w profile skewness were predominantly characterized by higher skewness in females in comparison to men, with an average of Cohen’s *d* = .2438 across all parcels that had higher skewness values for females than for men (*SD*_*all pos parcels*_ = 0.0700). These differences represent a more equal ratio of T1w/T2w signal intensity in superficial to deep cortical layers in females in comparison to men. The observed differences were predominantly located in transmodal areas, including the anterior cingulate cortex, insular areas, and the prefrontal cortex. Only the left medial occipital cortex presented the opposite pattern, such that the ratio of T1w/T2w signal intensity was more uniform in men, while the lower profile skewness values in females represent a stronger dominance of signal intensity in deeper layers in this area (mean effect size of all FDR-corrected negative effects *d* = -.3261, *SD*_*all neg parcels*_ = 0.0424).

Lastly, evaluating sex differences in the microstructural gradient, we found a significant shift towards both the lower and the upper extremes of the microstructural gradient for females in comparison to males (**Figure 2**A). In females, areas in red have a higher microstructural covariance with the gradient’s ‘upper’ anchor in ‘fugal’ (limbic and temporal) areas than men, and blue areas have a higher microstructural similarity with the gradient’s ‘lower’ anchor in sensory-motor and primary sensory areas in comparison to males (**Figure 2**A). Females’s left medial occipital areas as well as the right temporal pole were found to be more microstructurally similar to the sensory anchor of the microstructural gradient relative to males (mean effect size of all FDR-corrected negative effects *d* = -.2931, *SD*_*all neg parcels*_ = 0.0866). The bilateral supramarginal gyrus, parts of the inferior parietal cortex, and right anterior cingulate cortex were more similar to the ‘upper’ anchor in ‘fugal’ areas for females than for males (mean effect size of all FDR-corrected positive effects *d* = .2355; *SD*_*all* *pos parcels*_ = 0.0441).

We repeated all analyses additionally controlling cortical thickness as well as for family structure to account for potential confounds of twins in the dataset. Neither changed the original results (supplement 1).

We then tested the consistency of our results by quantifying the split-half-reliability for microstructural mean profiles, skewness and MPC gradient results (Churchill et al., 2013). We repeated our analysis in a 1,000 independent split-halves of our dataset (with equal ratios of males and females), and determined the mean spatial correlation for each of the three measures between the independent halves, respectively. The mean spatial correlation between the t-statistic maps of split-halves for profile mean was *r =* .8802 across the 1,000 permutations, the 5% 95% CI ranging between *r* = .8156 and .9227 across all 1,000 tests. The hippocampal values were similarly high, with a mean of *r* = .8093 (CI [5%, 95%] = .7481-.8549) for the left, and a mean of *r* = .8327 (CI [5%, 95%] = .7818-.8716) for the right hemisphere (for figure, see supplement 2). For profile skewness, split-halves spatial overlapped with a Pearson’s *r =* .7718 between t-statistic maps (CI [5%, 95%] = .7173-.8111). Only sex-differences in the gradient were less reliable, with a mean correlation of *r =* .6136 of t-statistic maps between split halves, ranging between *r =* .5445 and *r =* .6698. Overall, within this large cohort of healthy adults, observed sex differences in intracortical microstructure were thus highly to moderately reproducible.

### Sex differences in intracortical microstructure vary as a function of approximated sex hormone concentration (Fig 3.)

We hypothesized that sex hormones play a substantial role in shaping cortical microstructure. Hence, we expected that differences in menses-related hormonal profiles would influence the previously reported sex-differences. To test variation across hormonal status, we built five female subgroups that were determined by proxies for their current estrogen and progesterone concentration using a normative model of cyclic variations as well as by OC intake. We repeated the previous male vs. female contrasts five times, every time considering only those subgroups of females that were characterized by a certain hormonal profile: females who regularly took OC (n = 170), females who reported to be around their menstruation (low estrogen, n = 100); females who reported to be around their ovulation (high estrogen, n = 184); females who reported to be in their follicular phase (low progesterone, n = 171) and females who reported to be in their luteal phase (high progesterone, n = 113) (**Figure 3A**). Building on evidence about neuroendocrine plasticity effects on the short and medium term, we find that the observed differences between males and females vary as a function of menstrual cycle phase and regular OC intake.

**Figure 3.**
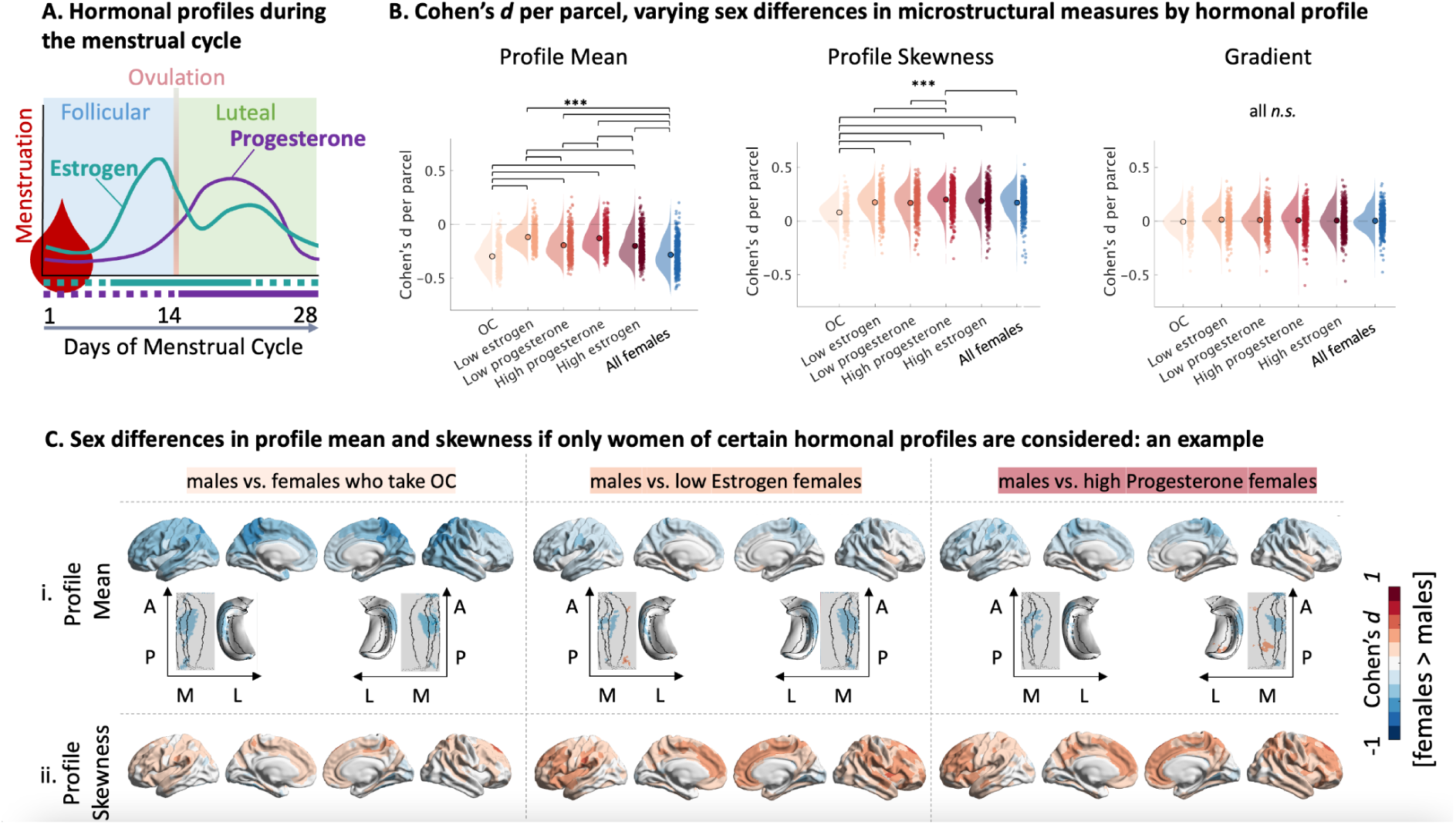
Comparing males to different female sub-samples, grouped by menstrual cycle phase. (A) Estrogen and progesterone fluctuate with the menstrual cycle. Horizontal lines under the x-axis indicate grouping: purple reflects progesterone (dotted = low; solid = high); turquoise reflects estrogen (dotted = low; solid = high) (B) Hormones determine cortex-wide sex-difference effect sizes based on post-hoc contrast on cortex-wide effect sizes. Cohen’s *d* per parcel is plotted separately for the three intracortical measures profile mean, profile skewness and the gradient, respectively for each sub-group-comparison. All shown contrasts were significant (p < .001). (C) FDR-thresholded Cohen’s *d* maps of T1w/T2w profile *mean* (i) between males and subsamples of females divided by OC use and menstrual cycle phase projected on the cortical surface and the hippocampus. (ii) FDR-thresholded Cohen’s *d* maps of T1w/T2w profile *skewness* between males and female subsamples mapped on the cortex.

When comparing effect sizes between different group comparisons for microstructural *profile mean,* only the OC-group could replicate the initial sex difference effect. Similarly to the previously reported effect, males had a significantly higher microstructural profile means in most parcels across the cortex when comparing them to females who regularly took OC (cortex-wide average *d*_*OC females* − *men*_= −0.2973). In contrast, sex difference estimations based on any other subgroup yielded significantly different cortex-wide effect sizes from the OC and initial group comparison (all p < .001, Figure 3B). This was especially evident for females who were estimated to have low estrogen or high progesterone levels at the time point of imaging (cortex-wide average *d*_*low estr females* − *males*_ = −0.1176; cortex-wide average *d*_ℎ*ig*ℎ *estr females* − *males*_ = −0.12846). The previously reported negative sex differences disappeared or even changed sign such that females presented a higher mean when comparing males to females estimated to have low estrogen or high progesterone levels (**Figure 3B, C**). We found that sex differences in the cingulate cortex, the insula, the orbitofrontal cortex and the hippocampus were most affected by the menstrual cycle phase and exogenous sex hormone intake (**Figure 3C**). For example, in the hippocampus, the higher *microstructural mean* in males’ compared to females’ medial CA2 and CA3 was the only sex difference that survived multiple comparisons when compared to females who took OC. In contrast, females medial CA2 and CA3 were much more similar to the males when females were in their high progesterone phase. The higher profile mean in the female CA1 in comparison to the male hippocampus was only visible bilaterally for females estimated to have high estrogen and low progesterone profiles (supplement 3).

Microstructural skewness varied slightly less as a function of cyclic hormonal variation. Here, The groups estimated to be in their low estrogen, their low progesterone and their high estrogen phase all replicated the initially reported sex difference in dominance of higher vs. lower cortical layers (all cortex-wide effect size contrasts *n.s.,* **Figure 3B**). However, the previously reported sex difference in microstructural profile skewness nearly disappeared when comparing males to females who regularly take OC (cortex-wide average *d*_*OC females*_= 0.0788, **Figure 3B**), and was even more pronounced when comparing males only to females estimated to have high progesterone concentrations (cortex-wide average *d*_ℎ*ig*ℎ *prog females*_= 0.1995). While the occipital areas, where the cortical microstructure in males was more uniform than that in females, were not affected by menstrual cycle phase, the change in the average effect size was particularly driven by stronger effects in the prefrontal, anterior cingulate and tempo-parietal areas (**Figure 3C**). In these areas, the signal intensity of lower and upper cortical layers was more uniform in females than men.

Comparing the microstructural gradient of males only to subgroups of females of different estimated hormonal profiles changed the distribution but not the mean of cortex-wide sex differences (all cortex-wide effect size contrasts between any group comparison *n.s,* Figure 3B).

Again, the sex difference effect varied strongest when comparing males to only OC takers versus comparing males to only females estimated to have high progesterone levels: Sex differences between OC takers and males were least extreme (min *d*_*OC females*_= -.4636, max *d*_*OC females*_= .3134), while sex differences between males and females in their high progesterone phase showed particularly big positive and negative effect sizes (min *d*_ℎ*ig*ℎ *prog females*_= -.5980, max *d*_ℎ*ig*ℎ *prog females*_= .3398). The posterior cingulate cortex, the dorsomedial prefrontal cortex and the medial prefrontal cortex all varied in how strong the shift to the ‘lower’ anchor in sensory-motor and primary sensory areas in microstructural similarity was in females of different hormonal profiles in comparison to men. Areas which varied with hormonal profiles in their positive sex effect, i.e. whose microstructural profiles was shifted more towards the ‘upper’, ‘fugal’ end of the gradient in females of different hormonal profiles in comparison to men, were the temporoparietal junction, the anterior cingulate cortex, the pregenual anterior cortex and the ventrolateral prefrontal cortex (see supplement 4).

### Endocrine plasticity effects on intracortical structure spatially overlap with cortical expression patterns of sex hormone related genes (Figure 4)

Since the sex differences we found varied strongly depending on which hormonal profile we approximated for females, we next asked whether the relevance of sex hormones in these sex difference effects could be supported on a molecular basis. A high density of sex hormone relevant genes in areas that express strong sex differences supports the notion that sex hormones play an important factor in microstructural sex differences. We thus next computed Spearman correlations between the transcriptomic maps of 25 sex steroid relevant genes and the sex-difference effect maps for each microstructural measure (Cohen’s *d* of sex differences in microstructural profile mean, skewness and covariance gradient).

**Figure 4.**
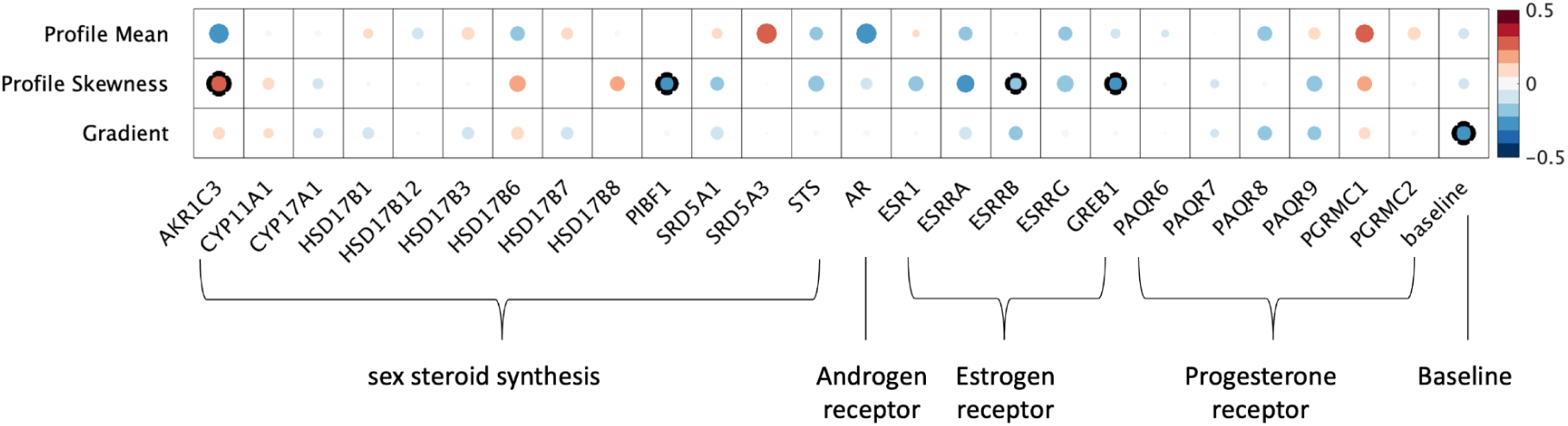
Spatial overlap between effect maps of sex differences for the microstructural gradient, profile mean and profile skewness. Transcriptomic maps of genes are sorted by categories: sex hormone synthesis related genes, androgen receptor related, estrogen receptor related genes, and progesterone receptor related genes. We test for spatial specificity by comparing against the principal component of all genes (baseline). Shades of red represent positive r-values, shades of blue represent negative correlations; circle size and shading indicate size of correlation. Significant p-values FWE corrected after spin-testing are marked with a black outline.

Sex-differences in *microstructural profile mean* were strongest associated with the transcriptomic map of the androgen-receptor activation related gene SRD5A3 (r = .31, *p*_*spin*_ = .06) and AKR1C3 (r = -.30, *p*_*spin*_ = .10). Strong overlap were additionally presented by the androgen receptor gene AR (r = -.31, *p* = .15) and the progesterone receptor PGRMC1 (r_*spin*_ = .26, *p*_*spin*_ = .20). These correlations, however, did not survive stringent correction for multiple comparisons. Sex-differences in T1w/T2w *microstructural profile skewness* demonstrated strong spatial associations with Progesterone Immunomodulatory Binding Factor 1 (PIBF1, r = -.25, *p*_*spin*_ < .05), the estrogen receptor beta (ESRB, -.22, *p*_*spin*_ < .05) and the Growth Regulating Estrogen Receptor Binding 1 (GREB1, r = -.24, *p*_*spin*_ < .05). While not surviving stringent multiple comparison control, all other estrogen receptor maps exhibited moderate negative associations with the microstructural skewness sex-effects (ESR1, r = -.18, *p*_*spin*_ = .10; ESRA, r = -.24, *p*_*spin*_ = .27; ESRRG, r = -.22, *p*_*spin*_ = .27). Lastly, sex differences in skewness also overlapped with the sex-hormone synthesis relevant gene AKR1C3 (r = .31, *p*_*spin*_ = .09).

The gene specificity for profile mean and the profile skewness sex difference was supported by a non-significant and negligible correlation with the baseline gene map we extracted. This was, however, not the case for the *microstructural gradient*, which correlated stronger with the baseline gene factor than with any other transcriptomic map (r = -.28, *p*_*spin*_ < .05).

### Sex differences differ in strength as a function of cytoarchitectural type (Figure 5)

Lastly, we sought to investigate the cytoarchitectural communalities of areas in which we had identified microstructural sex differences. Cytoarchitectural properties, such as laminar differentiation, are suggested to relate to plasticity (García-Cabezas et al., 2019). We thus tested if the sex difference effects contrived with the level of laminar differentiation by computing spatial correlations between effect maps of the identified sex differences and the hierarchy of *cortical types*. These cortical types, originally defined by von Economo and Koskinas (von Economo & Koskinas, 1925) and recently revised and histologically validated (García-Cabezas et al., 2020), describe a hierarchy of laminar differentiation from highly structured koniocortex to more diffusely structured agranular areas (Figure 5A)(García-Cabezas et al., 2019).

**Figure 5.**
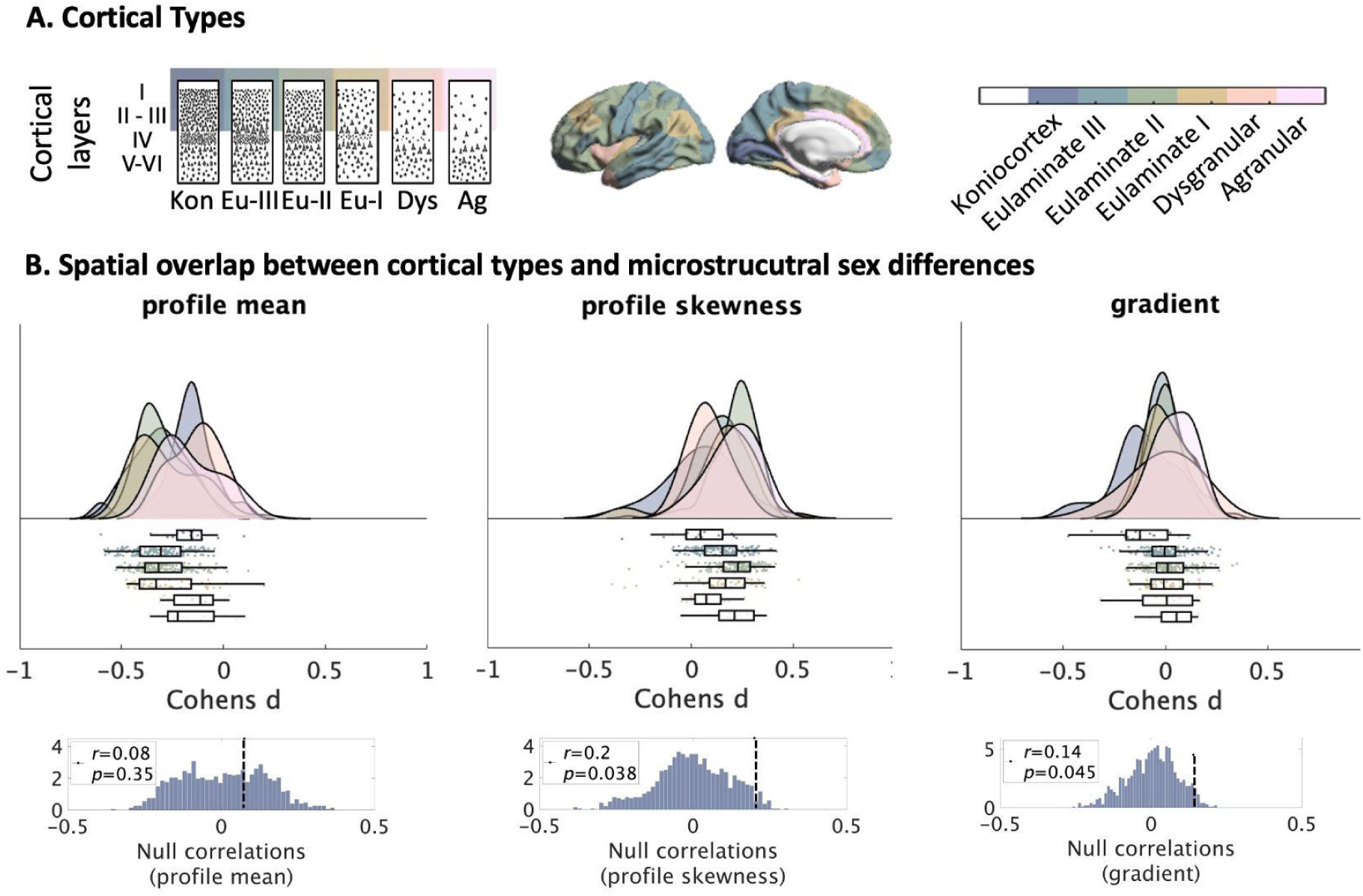
Contextualization of effects by histological decoding. (A) Schematic of Cortical Types according to von Economo and Koskinas and Garcia-Cabezas. (B) Results were put into context by spatial correlations with a hierarchy of laminar differentiation (cortical types). Figures show links between cortical type hierarchy and effect values (Cohen’s d per parcel) for each of the T1w/T2w profile-based intracortical measures. Raincloud plots show effect values per parcel, binned by cortical type. The bottom row visualizes zero-distributions between random hierarchies and effect maps in comparison to the statistical r-value.

We found that the effect size of sex differences in microstructural skewness and the microstructural gradient correlated with the hierarchy of cortical types, but not for microstructural mean (Figure 5B). A positive correlation between T1w/T2w profile skewness (r = .20, *p*_*spin*_ < .05) and cortical types indicated that differences in profile skewness were stronger for areas that can be characterized by less laminar differentiation (agranular mean Cohen’s d = .21, eulaminate cortex II mean Cohen’s d = .23). These results overlap with predictions about higher plasticity of these cortical types, and thus higher sensitivity to modulatory factors of plasticity such as sex hormones. Lastly, sex-difference effects in the microstructural gradient showed moderate overlap with the hierarchy of cortical types (r = .14, *p*_*spin*_ < 0.05). However, those regions with strongest microstructural gradient sex difference effects were in the most structured konio-cortical areas (mean Cohen’s d = -.13) and the most diffuse agranular areas (mean Cohen’s d = .06)

### Cerebrovascular control analyses

Changes in cerebrovascular blood flow could pose a potential confound that explains microstructural variation with the menstrual cycle. In addition to including intracranial volume as a covariate in every linear model, we thus tested if the relation to sex hormone concentration would covary with the local density of cerebral vasculature.

We found that sex-differencers in T1w/T2w profile mean overlap moderately with cerebral vein density (*r* = .28, *p*_*spin*_ = .065). This overlap is stronger for sex differences in profile mean comparing males to ovulating female sub-groups: profile mean differences in males compared to females in their low progesterone phase (*r* = .28, *p*_*spin*_ = .05), in males compared to females in their low estrogen phase (*r* = .29, *p*_*spin*_ < .05), in males compared to females in their high estrogen phase (*r* = .32, *p*_*spin*_ < .05) and in males compared to females in their high progesterone phase (*r* = .35, *p*_*spin*_ < .05), can be significantly associated with cerebral vein density.

No sex difference effect in T1w/T2w profile skewness overlapped significantly with any cerebral vasculature atlas (all *n.s.*).

Lastly, while differences in the microstructural gradient between males and the collapsed female group did not correlate with the arteries (*r* = .07, *p*_*spin*_ = .156) or veins (*r* = -.11, *p*_*spin*_ = .162) atlas, they did for some female subgroups. The effect maps showed greater overlap with cerebral artery density (*r* = .17, *p*_*spin*_ < .05) and in cerebral vein density (*r* = -.21, *p*_*spin*_ < .05), if only females in their high progesterone phase were considered when looking at sex differences in the microstructural gradient.

## Discussion

A crucial element in understanding how brain structure supports brain function is grasping the dynamics of brain structure: structural plasticity. Nevertheless, most brain structure studies fail to take major cortical plasticity principles into account, such as neuroendocrine plasticity. Neuroendocrine plasticity systematically shapes brain structure through molecular signaling cascades, being the major source of sex differences in brain structure. While a large amount of neuroimaging studies have reported the effect of sex hormones on macro-level brain structure, the microstructural changes that underpin these variations are not well understood. Here, we thus aimed at highlighting associations between sex differences in cortical microstructure and sex hormones. The present study set out to characterize systematic variations in microstructural measures between males and females, and evidence how these are related to gonadal hormones. We approximated hormone concentration from menstrual cycle data, and investigated the spatial specificity of sex hormone related genes for the observed sex differences. Making use of T1w/T2w signal profiling within the cortical sheath, we derived an average measure of intracortical tissue properties (microstructural profile mean), a measure of the distribution of layers within the cortex (microstructural profile skewness), and a measure of the relative distribution of microstructural organization across the cortex (principal microstructural gradient). Using these microstructural measures, we first examined sex differences in a large cohort, and then used a multi-modal approach to identify neuroendocrine correlates of these identified differences, with particular focus on sex hormones. We found that the cortical microstructure of males and females differ regionally in each of these microstructural measures. Importantly, the effect size of the observed sex-differences depended on the estimated estrogen and progesterone levels of females at the time of the brain scan, and spatially overlapped with expression levels of sex-hormone-relevant genes. This was strongest for the cytoarchitectonic proxy measure of laminar differentiation, where sex-differences mainly appeared in dys- and agranular areas. These types of less structured cortex have been suggested to display comparably strong effects. We statistically controlled for ICV in all models and provide evidence that the observed effect was not confounded with hormone-induced fluctuations in cerebrovascular blood flow.

### Systematic variation in cortical microstructure between males and females

Sex hormones influence brain structure most strongly during sensitive periods of development, such as the perinatal period (Juraska et al., 2013) and puberty (Sisk & Zehr, 2005), but continue to shape brain structure over the lifetime. Here, we computed a snapshot of this long-term neuroendocrine plasticity effect by contrasting the cortical microstructure of adult males and females and then associated these effects with sex hormones in two complementary analyses.

The male cortex was characterized by an overall higher T1w/T2w signal mean spreading the whole cortex, except for bilateral insular and medial temporal areas. The microstructural profile mean reflects intracortical myelin (Glasser et al., 2014; Glasser & Van Essen, 2011), as well as iron concentration (Uddin et al., 2016), cell density and water content. This measure thus describes differences and similarities between a combination of intracortical tissue properties. Myelin (Darling & Daniel, 2019; Glasser & Van Essen, 2011), iron concentration (Anttila et al., 1997), cell density (Y. Kim et al., 2017) and water content (Lorio et al., 2016; Schulte-Hostedde et al., 2001; Stüber et al., 2014) are all subject to sexual differentiation triggered by sex hormones especially during critical periods of development. For example, sex hormones released during puberty lead to myelination sex differences in the rat prefrontal cortex (Darling & Daniel, 2019). The widely spread sex difference effect in our average measure of intracortical tissue properties confirms this notion. On the other hand, the combination molecules that influence mean T1w/T2w also make it the most prone to confounds, such as transmit bias field effects (Glasser et al., 2022), and sex hormone effects on cerebral fluids. The moderate overlap mean T1w/T2w effects with cerebral vein density furthermore might be an interaction with the effect of venous blood on T_2_w signals (Sedlacik et al., 2008). Since profile skewness and the microstructural gradient are based on relative variations of T1wT2w, they do not suffer from the same limitations. Together, our results might both be explained by a stronger cortical myelination in males or female hormonal effects on cerebral fluids. Histological studies are required to disentangle the two.

Previous studies report hormone-related volumetric gray matter differences in regions including the precuneus, insula, ACC, the middle temporal lobe, inferior frontal gyrus, middle frontal gyrus, superior frontal cortex, and hippocampus (W. Qiu et al., 2004; Rehbein et al., 2021). Here, we add a more nuanced characterisation of these differences by showing that females in comparison to males show a more skewed microstructural pattern in temporo-parietal, precuneus, insular and frontal areas. Phrased differently, these areas exhibit a comparatively less pronounced differentiation between supra and infragranular cortical layers in females compared to men. Only in the left medial occipital cortex does this trend invert, revealing a heightened dominance of signal intensity in deeper layers in females’s cortex.

We observed less pronounced but similar findings using a gradient approach, demonstrating that bilateral temporal-parietal regions are characterized by higher gradient loadings, e.g. are more similar to fugal anchors, in females relative to men, yet visual areas are more similar to the sensory anchor in females relative to men. The occipital lobe is a koniocortical area characterized by six clearly distinguishable cortical layers (Fatterpekar et al., 2002). Our findings indicate that the clear cytoarchitectural differentiation in this area is stronger in females in comparison to men. Men’s occipital lobe, comparably, has more cytoarchitectural similarities with less clearly structured cortical types. Reversely, the temporo-parietal junction and medial sensory-motor areas in females are more similar to areas that are typically characterized by less structured cytoarchitecture. Overall, the covariance of microstructural profiles thus shifted more towards the extremes in females, while males exhibited a more gradual change of cytoarchitectural profiles. The overlap of significant areas with our skewness results shows that this can be mainly explained by differences in profile skewness, indicating a stronger differentiation between upper and deeper cortical features in females relative to males in terms of microstructure. Notably, while the overlap of these findings validate the measures respectively, it is important to note that our reliability measure was least consistent for the gradient approach.

Overall, the regional distribution of sex differences in microstructural measures overlaps with previous reports on sex differences in gray matter volume in the same cohort of participants, in particular in cingulate and frontal areas (Liu et al., 2020). Moreover, temporal-parietal, frontal and insular regions are also found to display a diverging coupling of structure and function between sexes (Gu et al., 2021). Indeed, in related work in the same sample (Valk et al., 2022), our group observed increased coupling of function and microstructure in females in regions that show heightened skewness in females. Follow-up studies that focus on the functional implications of the reported microstructural measures are required to shine light on functional implications of the reported microstructural sex differences.

### Evidence for the influence of sex hormones on microstructural variation between males and females

Many factors contribute to diverging brain structure between sexes. Typically, a complex interaction between genetic, epigenetic, gonadal and hormonal factors give rise to differences in brain structure (Grgurevic & Majdic, 2016; McCarthy & Arnold, 2011). Acknowledging this complexity, here we specifically investigated whether sex hormones could be associated with the observed sex differences in microstructure. We followed up our initial sex difference analysis with two orthogonal hormonal analyses. First, we systematically compared if sex differences would change if female subjects were grouped by different approximated hormonal profiles. We focus on hormone concentration throughout the menstrual cycle, which changes up to 80 fold (Mihm et al., 2011; Stricker et al., 2006), as well as on regular OC use. The synthetic estradiol and progesterone of OC bind to estrogen and progesterone receptors, which leads to overall flattened and lower endogenous estrogen (Kjeld et al., 1976) and progesterone (Rabe et al., 1997) levels via negative feedback mechanisms (Baird & Glasier, 1993; Frye, 2006; Porcu et al., 2019; Rivera et al., 1999). Regular OC-intake thus changes females’ hormonal profiles on a medium term.

### Microstructural sex difference effects change with menstrual cycle phase and oral contraceptive use

In line with evidence of macrostructural short-term hormonal plasticity studies (De Bondt et al., 2013; Dubol et al., 2021; Lisofsky et al., 2015; Ossewaarde et al., 2013; Petersen et al., 2021; Pletzer et al., 2010; Rehbein et al., 2021; Romeo et al., 2004), also our microstructural measures changed when comparing males to groups of females with different estimated hormonal profiles, yielding the first piece of evidence that sex hormones play a role in microstructural sex differences in the human cortex. Areas in which the effect size or even the directionality of the sex difference effect varied strongest depending on the hormonal profile of the females, largely overlapped with regions that had previously been named as key regions for volumetric menstrual cycle differences (hippocampus, cingulate cortex, insula, inferior parietal lobule, prefrontal cortex (Dubol et al., 2021)), or gray matter volume differences due to oral contraceptive use (prefrontal cortex (De Bondt et al., 2016) and the cingulate cortex (De Bondt et al., 2013)). Adding to these observations of the effect of sex hormones on macro-level brain structure, our result contributes with a more nuanced understanding of microstructural property variations.

For the average cortical microstructure profiles, we could only replicate the sex difference effect with females who took oral contraceptives. This outcome underscores the potential influence of exogenous hormone manipulation on the cortical microstructure and highlights the robustness of the sex difference effect observed in this subgroup. Comparisons to all other subgroups significantly changed the sex difference effect for this microstructural measure. Indeed, previous work indicates an interplay between glia and sex steroids, impacting myelin formation and organization (Garcia-Segura & Melcangi, 2006). The sex difference effect varied most notably with sex hormones in the cingulate cortex, the insula, the orbitofrontal cortex and the hippocampus. For example, the average profile difference in the CA2 and CA2 subfields, which was much higher in males than females in the overall sample, was non-significant when only comparing males to females at their highest progesterone levels. This hippocampal finding adds a more detailed understanding to recent findings which link estrogen and progesterone variations specifically to macro-scale structural changes in the hippocampus (Taylor et al., 2020; Zsido et al., 2023).

In contrast to the mean microstructural profiles, where effect sizes exhibited more substantial variations across different hormonal subgroups, microstructural skewness displayed a relatively stable pattern. The low estrogen, low progesterone, and high estrogen groups all replicated the initial sex difference in the dominance of higher versus lower cortical layers. However, the effects were significantly different when examining females who regularly took oral contraceptives or had high progesterone concentrations. Specifically, there was nearly no difference in lamination between males and females who took OC, but there was an even stronger difference in lamination between males and females with high progesterone concentrations. These variations were mainly driven by stronger effects in the prefrontal, anterior cingulate and tempo-parietal areas. This suggests that in the case of microstructural layer variations, the initial sex difference effect was driven by naturally cycling females, and that medium-term effects of oral contraceptives specifically contribute to a reduction or exacerbation of cortical layer dominance, making this microstructural feature in females more similar to men. Similar to the main sex difference result, for hormonal variations, the microstructural covariation pattern again replicated the skewness finding. Strongest deviations from the initial sex differences were found for females who took OC, and for females in their high progesterone phase, in the cingulate cortex (posterior and anterior and pregenual anterior cortex), several prefrontal areas (dorsomedial prefrontal cortex, medial prefrontal cortex, ventrolateral prefrontal cortex) and the temporoparietal junction.

It is important to note that rather than longitudinally following up on microstructural changes going along with hormonal variations intra-individually, here we computed inter-individual contrasts on an indirectly approximated correlative hormonal measure. We thus merely interpret our results as tendencies which underline the importance of considering the complexity of hormones in the study of brain structure. However, since we benefit from a big sample size and a second, independent hormonal analysis, our results underscore the importance of moving beyond a generalized understanding of sex differences and considering hormonal profiles as a crucial factor in interpreting and explaining these differences.

### Cortical microstructure differs in areas with rich sex hormone related gene expression levels

To support the evidence of our first endocrine analysis, we added a second, independent one. We show that cortical microstructural differences that we systematically observe between males and females are specific to areas with elevated expression levels of sex hormone related genes. This offers a translation of a recent rodent study to humans, where sex differences in brain structure occurred particularly in regions enriched in sex hormone genes (L. R. Qiu et al., 2018). This orthogonal hormonal analysis yields the second piece of evidence that sex hormones contribute to microstructural differences in the human cortex.

Regions in which the microstructural profile mean was higher in males than females are rich in androgen receptors (AR) and regions where this average microstructure measure is higher in females than men, strongly express the membrane-associated progesterone receptor (PGRMC1). Both the androgen receptor (Bielecki et al., 2016) and progesterone receptors (Ghoumari et al., 2020) have a key role in myelination. Progesterone and its metabolites interact with oligodendrial differentiation and thus affect development of oligodendrocytes and myelination (Gago et al., 2001). Since microstructural profile mean (at least in parts) reflects intracortical myelin levels and due to limitations of neuroimaging, we hypothesize that myelination rather than synaptic plasticity effects or dendritic remodeling could drive the results observed in this study. Future studies observing causal links on a molecular level are needed to confirm this hypothesis. In contrast, while sex differences in the average cortical microstructure measure tended to overlap with myelin-plasticity-related genes, systematic differences in microstructural lamination rather overlapped with expression levels of genes that were linked to neural plasticity mechanisms. For example, the estrogen receptor genes are implicated in glutaminergic synapse formation (Jelks et al., 2007), neurogenesis, synaptic spine density (M. Kim et al., 2019), synaptic plasticity (Bäumler et al., 2019) and neural differentiation (Pillerová et al., 2021). Lastly, genes important in the metabolism and thus supply of sex hormones such as AKR1C3 (W. Qiu et al., 2004, p. 20), PIBF1(Liu et al., 2020) and SRD5A3 (Uemura et al., 2007) overlapped both with sex differences in cytoarchitectural lamination and the mean cytoarchitectural measure. Together, this multitude of possible plasticity effects linked to the observed microstructural differences suggests a complex molecular interaction rather than a linear causal chain in the role of sex hormones in cortical microstructure. To support this claim and go beyond the correlative nature of this result, causal cytoarchitectural studies which include genetic manipulation are required.

### Sex hormones in the hippocampus

Numerous hormone-related neuroimaging studies finds the hippocampus to be affected by sex-hormone induces plasticity(Lisofsky et al., 2015; Pletzer et al., 2018; Protopopescu et al., 2008; Zsido et al., 2023). We thus made efforts to extend our cortical-surface based analysis to the hippocampus by projecting an average measure of cortical microstructure on this unfolded surface. As expected, we found marked sex-differences that differed as a function of hippocampal subfield, with subiculum showing higher T1wT2w values in females, but CA2/3 showing higher T1wT2w mean in males relative to females. Though research on sex differences in hippocampal structure in humans mainly focuses on the whole hippocampal volume, recent work has indicated marked changes in particular in subicular microstructure (Bayrak et al., 2022) and volume (van Eijk et al., 2020). Previous work in rats has shown marked changes in CA1 and CA3 but not CA2 (Yagi et al., 2020). Such differences between sexes were mainly attributed to hormone-related effects. Indeed, when evaluating effects of sex hormones on hippocampal structure, we observed that the relatively stronger T1wT2w signal in females in the subiculum wasn’t present when comparing males to females taking oral contraceptives or females in their low estrogen phase. Conversely, these subgroups showed even lower average T1wT2w values in CA subregions relative to men. A recent longitudinal study furthermore found that CA1 volume decreases for high progesterone concentration in the menstrual cycle (Zsido et al., 2023). Here, we complement this finding and show that sex differences in this area nearly disappear when females are in this part of their menstrual cycle. Overall, these findings extend previous work showing an association between hippocampal microstructure and sex hormones. Through unfolding the hippocampus we increased regional specificity, taking into account the morphology of the hippocampus (DeKraker et al., 2020). Further work studying the impact of sex hormones on hippocampal structure may use similar techniques to capture regional variation.

### Cortical cytoarchitectural differentiation supports endocrine plasticity and stability

Lastly, given our technique was heavily inspired by traditional neuro-anatomy procedures(*Brodmann: Die Rindenfelder Der Niederen Affen - Google Scholar*, n.d.; Palomero-Gallagher & Zilles, 2019; von Economo & Koskinas, 1925), we aimed to embed our results within current cytoarchitectural models. Demands to dynamically adapt brain structure vary across the whole brain: for example, it is adaptive that adult sensory areas respond in the same way to the same sensory inputs, while higher-order areas need to flexibly adjust their ways of processing depending on previous life experiences (Mesulam, 1998; Valk et al., 2020, 2022, 2023). Consequently, stability of neural structures is adaptive in sensory cortices, while plasticity is adaptive in higher order structures.

This divergence between the need of plasticity and stability covaries with cortical microstructure. Sensory input to the brain is perceived and processed in idiotypic and unimodal areas, which are the cytoarchitectonically most elaborate areas (highly structured koniocortex and eulaminate cortex III). The paralimbic cortex, on the other hand, receives projections both from other cortical areas-mainly from higher-order sensory and association cortices, such as the auditory association cortex, temporal sensory association areas and other prefrontal cortices (Barbas, 1995; Barbas et al., 1999), as well as from subcortical structures, such as the amygdala and the thalamus (Dermon & Barbas, 1994). Supporting the hypothesis of a higher need of plasticity in areas that are not directly linked to sensory or motor organs, plasticity has been shown to systematically vary with laminar elaboration (García-Cabezas et al., 2017), so that less elaborate structures are the most *plastic*, and most elaborate structures present the most *stability* markers.

Here we show that this gradient from plastic to static cytoarchitecture extends to endocrine plasticity, specifically for cortical laminar differentiation. The more plastic the cortical type of a brain area, the stronger the effect of sex hormones we observed for this measure. Apart from sex hormone triggered second-messenger cascades, other molecular factors additionally support the plasticity of these cortical types. For example, they are richer in calmodulin-dependent protein kinase II (CaMKII), which is an enzyme known to be crucial for synaptic plasticity. Areas of higher granulation are richer in parvalbumin neurons, which limit plasticity via perisomatic inhibition of neighboring pyramidal neurons (García-Cabezas et al., 2017). Further, dendritic spine pruning is reduced in the adult limbic cortex in comparison to eulaminate areas, supporting LTP and synapse formation (Sasaki et al., 2015). Together, these cytoarchitectural properties paired with the appropriate hormone receptor infrastructure may allow the cortex to adapt readily to varying demands commanded by fluctuating levels of sex hormones over short, medium and long time scales.

### Cerebrovascular control analyses shows overlap with T1w/T2w signal mean, but not others

Work in the field of endocrine plasticity has been critiqued in the past for its potential confounds with hormone-related blood flow changes and water shifts in the brain. Since the MR signal and in particular T1wT2w contrast (Glasser et al., 2022) is both sensitive to myelin, but also to water and iron, as well as impacted by the radiofrequency transmit field (B1+), it is not straight-forward which effects we determined with our analyses. To limit these uncertainties, we conducted two control analyses: first, we included intracranial volume as a covariate in each of our linear models, statistically controlling for hormone-induced volume fluctuations. Second, we quantified the overlap between cerebral vasculature and the areas in which identified sex difference effects. The correlation with hormonal mean T1w/T2w profile, but not skewness supports the notion that T1w/T2w signal may indeed be globally modulated by the effect that sex hormones exert on water-balance and lipid metabolism. Third, conceptually intra-cortical metrics may be least biased by these features as they reflect a relative metric perpendicular to the cortical sheet, implicitly correcting for spatial variations across the cortex.

## Conclusion

In this study, we performed multi-level analysis on sex differences and the influence of sex hormones on the average measure of cortical microstructure, laminar differentiation within the cerebral cortex and the microstructural gradient. Effects on laminar differentiation were consistent across the sample, varied systematically between hormonal subgroups, and strongly overlapped with sex hormone gene expression levels. Adding to this, we found that this measure was not affected by vasculature. Together, our study advances our understanding of how cortical microstructure relates to sex differences and hormonal variations. The results emphasize the need to consider hormonal subgroups when investigating such differences, as these subgroups can yield diverse and sometimes contradictory outcomes. Moving forward, continued research in this area will contribute to a more comprehensive comprehension of the interplay between sex differences, brain structure, and hormones, ultimately enhancing our insights into the intricate underpinnings of human brain function and behavior.

## NOTE FROM THE AUTHORS [societal relevance and on politics in neuroscience]

The organization of brain structure is largely shared between sexes (Eliot et al., 2021). For those brain structures in which we find systematic differences, it is oversimplified to call ‘biological sex’ the cause. Sexual differentiation is complex, where several factors play a role. Amongst others, sex chromosomes, gonadal hormones, and epigenetic factors are the reason for systematically observed differences between men and women (Grgurevic & Majdic, 2016; McCarthy & Arnold, 2011). Because genes, gene expression and environment influence each other over the lifetime, there is no such thing as a ‘male’ or ‘female’ brain, but instead, a combination of several dynamic variables that influence brain structure. If the research on sex differences is aimed at serving a higher objective such as making drugs equally effective for men and women, it is thus crucial to contextualize the observed effects. In the present study, we therefore investigated the temporal stability and link between observed sex differences in intracortical cytoarchitecture and sex hormones and sex hormone receptor expression. In vertebrates, hormone receptor signaling may have evolved to coordinate gene regulation throughout a neural circuit as a strategy for controlling context-dependent behavioral states. Moreover, the association between hormone receptor target genes identified here and human neurological and neurodevelopmental conditions may explain the notable sex biases of these diseases.

## Methods

### Participants and study design

In this study, we leveraged the HCP S1200 young adult data release 147 (Van Essen et al., 2013). In the following, we reiterate the most important details of this work, but more details of the HCP study design are described elsewhere (Van Essen et al., 2013). The HCP dataset includes functional and structural MRI data acquired with 3T scanners from a total of 1206 healthy adult twins and their non-twin siblings born in Missouri, as well as behavioral and cognitive measures and extensive demographic and health-related data. Participants were recruited based on data from the Missouri Department of Health and Senior Services Bureau of Vital Records. The HCP consortium aimed at collecting a representative sample in respect to behavioral, ethnic, and socioeconomic diversity. To allow for sufficient variability in the healthy sample, only severe neurodevelopmental, neuropsychiatric and cardiovascular illnesses were excluded. For the present study, we only used structural MRI data and removed all subjects with missing MRI values, so that we included n = 1093 individuals (n = 298 monozygotic, n = 188 dizygotic twins, n = 449 not related individuals), out of which n = 594 were female. We classified those individuals as females who reported a female gender and are or have been menstruating in their lives. The age mean +- SD was 28.8 +- 3.7 years (age range = 22-37 years). The current research complies with all relevant ethical regulations as set by The Independent Research Ethics Committee at the Medical Faculty of the Heinrich-Heine-University of Duesseldorf (study number 2018-317).

### Neuroimaging Data Acquisition and Preprocessing

We obtained readily preprocessed T1-weighted (T1w) and T2-weighted (T2w) structural scans in 0.7 mm isotropic resolution from the HCP openly available dataset. MRI data used in this study were acquired with Siemens Skyra 3 T scanners (32 channels) customized for the HCP. Two T1w and two T2w images were collected in a total of 32 minutes, using identical parameters respectively. T1w was acquired with the 3D MPRAGE sequence (Mugler III & Brookeman, 1990) in 256 sagittal slices with an echo time of 2.14ms, an inversion time of 1000ms, and a repetition time of 2400ms (flip angle = 8°; matrix = 320). The T2w images with identical geometry as the T1w ones were acquired with the turbo spin-echo sequence (Mugler et al., 2000) allowing for variable flip angles, with an echo time of 565 ms, a repetition time of 3200 ms, and a bandwidth of 744 Hz per pixel. Data was preprocessed with the Freesurfer version 5.3. Amongst other steps, T1w and T2w images were aligned, corrected for field bias, segmented and their ratio (T1w/T2w) was projected to the cortical surface in FSaverage5. Detailed pipelines and preprocessing steps are described in (Glasser et al., 2013).

### Intracortical microstructure profiling & parcellation

In traditional neuroanatomy, intracortical depth-dependent measures are commonly used to describe micro-architectural characteristics of brain regions (Palomero-Gallagher & Zilles, 2019; Pijnenburg et al., 2021; Zilles et al., 2002). Analogous to this approach, cross-sectional profiles of T1w/T2w MRI signal intensity across the cortical mantle build the basis for several local and global estimates of different cytoarchitectural properties (Paquola, Wael, et al., 2019; Schleicher et al., 1999): First, the *mean* of T1w/T2w profiles perpendicular across the cortical mantle reflects the local mean T1w/T2w signal intensity across the cortical gray matter. Second, the *skewness* of T1w/T2w signal intensity across cortical depths contrasts local dominance of superficial to deep cortical layers (Zilles et al., 2002). Thus, microstructural skewness yields a proxy of laminar differentiation relating to cytoarchitectural complexity. In addition to these two *regional* measures of cortical microstructure, microstructural profile covariance (MPC) quantifies *global* variation of inter-regional microstructural similarity across the cortex (Paquola, Wael, et al., 2019). The utility of this approach has been demonstrated in previous studies that showed a gradient of microstructural differentiation is mirroring brain function and orchestrates brain development in adolescence (Paquola, Bethlehem, et al., 2019), and has been validated by comparing the MRI measure with traditional microscopy-based profiles (Paquola, Wael, et al., 2019). Using the two local metrics of microstructural profile mean and skewness, as well as the global metric of the microstructural profile covariance gradient thus allow to analyze different biologically relevant aspects of intracortical microstructure in vivo.

To build these measures from the preprocessed T1w/T2w surface data, we first computed 12 equivolumetric surfaces between the pial and white matter surface, generated by FreeSurfer. To compensate for cortical folding, the model varies the Euclidean distance ρ between two intracortical surfaces and thus preserves the fractional volume between them. The following formula was used to calculate ρ (A_out_ = outer cortical surface, A_in_ = inner cortical surface, α = fraction of the total volume of the segment accounted for by the surface).

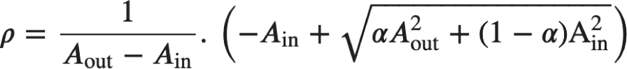

We sampled signal intensities along all linked vertices from the pial to the white matter surface across the whole cortex. To reduce computational efforts and multiple testing problems while accounting for a biological relevant heterogeneity and spatial specificity, we parcellated the data into 400 Schaefer parcels by computing the average value of T1w/T2w signal intensity per sample point for each of the 400 parcels (Schaefer et al., 2018). The Schaefer parcellation approach is based on resting-state functional MRI (rs-fMRI), using a gradient-weighted Markov Random Field model which takes local transitions in rs-fMRI patterns as well as similarity of global rs-fMRI patterns into account to define the parcels.

### Analysis of microstructural profiles across the whole sample: mean, skewness, and microstructural gradient

From the resulting 400×x12 data matrix (12 sample points across intracortical depth), we computed the microstructural profile mean and skewness separately for each parcel for each subject. For better interpretability, the skewness values were then rescaled to values between 0 and 1.

To extract the principal gradient of microstructural covariation in the cortex, we first built a MPC matrix by pairwise Pearson product-moment correlations, taking the average whole-cortex intensity profile into account. MPC was thus defined for each microstructural profile pair i, j and each participant s as:

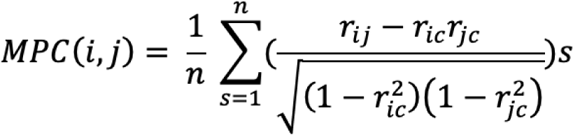

where r_ic_ is the correlation coefficient of the intensity profile at parcel i with the average intensity profile across the entire cortex, r_jc_ is the correlation between the intensity profile at each parcel j with the average intensity profile across the cortex, and n is the number of participants. Finally, we log-transformed and thresholded the MPC above zero and then only kept the top 10% of strongest microstructural similarity pairs.

We decomposed the MPC matrix (400 x 400 parcels) onto its low-dimensional representations by implementing the diffusion map embedding algorithm (Coifman & Lafon, 2006) using the BrainSpace toolbox (Vos de Wael et al., 2020). First, we calculated an affinity matrix of the MPC with a normalized angle kernel function and then decomposed it nonlinearly onto a set of 10 principal eigenvectors, namely the *gradients* (Margulies et al., 2016; Paquola, Wael, et al., 2019). In this gradient space, parcels that have similar microstructural profiles are situated closely to each other, whereas parcels that have distinct profiles fall apart. For each participant, gradients of MPC were obtained separately. However, to increase comparability across gradients, we then aligned the individual gradients with the gradient derived from the group-average MPC. All individual gradients were then rescaled to values between 0 and 1.

We repeated the analysis of all three measures regressing out cortical thickness and including the family structure (interaction between zychosity and family status) as a random effect to demonstrate that our results were not affected by these variables (supplement 1).

### Hippocampal unfolding & projection

The hippocampus is both a highly plastic brain structure and a structure with high density of sex hormone receptors and thus is a region of interest for our analysis. It is, however, not included in the freesurfer cortical projections. To nevertheless include this crucial brain structure in our analysis, we used an automated hippocampal segmentation pipeline - *HippUnfold* (DeKraker et al., 2022)-that projects hippocampal MRI values to a 2D surface while preserving its topological structure, similar to cortical surface projections. Since this procedure is more sensitive to poor data quality, we performed a more stringent quality assessment. Out of the initial 1206 subjects, we excluded n’ = 160 subjects with anatomical anomalies or tissue segmentation errors, n’ = 93 subjects for which no preprocess T1w images were available, and n’ = 86 with morphological outliers (thickness, surface area, gyrification, curvature or T1w/T2w values exceeding 2.5STD of group average), such that we overall included n = 867 subjects (n = 500 female, n = 367 male) in the hippocampal analysis. The pipeline for the surface projection is described in detail elsewhere (DeKraker et al., 2018, 2020). In short, the hippocampal regions of interest were first cropped from the preprocessed T1w/T2w data and the hippocampal ‘cortical surface’ was further segmented with a U-Net neural network architecture (Isensee et al., 2021). By solving Laplace’s equation, HippUnfold then transforms the segmented MR data from Cartesian native space to unfolded space. The transformed data was then stored in GIFTI files, so that any following analyses could take place as if it were surface data. However, due to the thin subregions and the complicated folding of the hippocampus, the previously described procedure of computing equivolumetric surfaces between outer and inner layer was not possible for the hippocampus. Instead, only the mean T1w/T2w ratio was used as a hippocampal MR measure, yielding one instead of three microstructural measures for the hippocampus.

### Sex-difference and proxies for links to sex hormones

We used different estimates of sex hormones to determine neuroendocrine plasticity in the short, medium, and long term. As a proxy for long-term effects of sex hormones, we first computed sex differences for each of the three T1w/T2w measures. We estimated sex differences with linear mixed effect models (LME) using the Matlab module of SurfStat (Worsley et al., 2009). In each model, we accounted for age and intracranial volume (ICV):

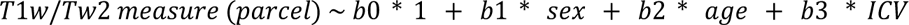

We then computed the t-statistics for the contrast ‘females - males’ for each of our three microstructural measures. LMEs were estimated separately for each parcel, and t-statistics were projected back to the cortical surface. We then two-sidedly corrected the t-values for a false discovery rate (FDR) of .05 (Benjamini et al., 2005; Benjamini & Hochberg, 1995). For easier comparison between tests, we report the effect size quantified by Cohen’s *d* for all results that reached significance after this defined FDR-correction threshold.

Secondly, we approximated hormone concentration levels in females to estimate neuroendocrine plasticity in the medium or short term. We used self-reported days since menstruation and about OC intake as a grouping variable. First, we subdivided females into two groups, those who regularly took OC (n = 170) and naturally cycling females (NC). Lastly, we built groups in which females differed in their hormonal profile on a short term. Since estrogen and progesterone concentration peak at different points within the menstrual cycle, we subdivided NC females in a low and high progesterone, and in a low and high estrogen group, respectively. In total, we thus compared five subsamples of females against the cortical microstructure of men: an OC group, a high and low estrogen group, and a high and low progesterone group. We included all females who reported regular menstrual cycles, and that their last menses was between 0 and 28 days (n = 284), which is considered the length of a normal menstrual cycle (Mihm et al., 2011). Estrogen is low in the beginning of the cycle and starts to rise before ovulation, with a second peak premenstrually in the luteal phase, before it drops again just before and during menstruation (figure 3 A). We accordingly built a high estrogen group for females who were broadly around ovulation (between day 7 until day 23, n = 284), and a low estrogen group for females that were just before and during menstruation (n = 100). Progesterone surges after ovulation during the luteal phase, and was thus defined as low before day 15 (n = 171), and high after day 14 (n = 113). This classification is in accordance with common comparisons between ovulation and menstrual (high and low estrogen) and luteal vs. follicular phase (high and low progesterone) (Frank et al., 2010; Lisofsky et al., 2015; Pletzer et al., 2010; Toffoletto et al., 2014). Based on these groups, we then modeled the three microstructural measures with five LMEs, in which we included all males together with one respective female subsample. As before, we included age, sex, and ICV as covariates, and computed a contrast for females - males for each of the five models. Because we were interested in comparing the effect sizes of each group-comparison with each other but sample sizes differed, we computed Cohen’s d for each parcel that survived FDR correction for multiple comparisons. Lastly, to determine whether effect sizes differed per group comparison, we computed a one-way ANOVA on the Cohen’s d values across the 400 parcels between groups and post-hoc contrasts based on Tukey’s honestly significant difference procedure.

### Genetic decoding

To complement our macro-level analysis of the effect of sex hormones on cortical microstructure on a molecular level, we then tested if sex hormone linked gene expression patterns would overlap with the observed effects. We selected genes of interest (GOIs) via open ontologies such as KEGG (https://rest.kegg.jp/get/hsa00140) and genecards (https://www.genecards.org/). We included genes that were either relevant in the *synthesis process* of the standard sex hormones (testosterone, estrogen, progesterone, adrenal androgens: dehydroepiandrosterone and androstenedione, progesterone-derived neurosteroids: allopregnanolone and pregnenolone), or linked to androgen, estrogen or progesterone *receptors*. In the end, we included n = 25 GOIs for which we had access to transcriptomic expression maps on the cerebral surface (see supplementary table 5). Our list of GOIs largely overlapped with previous selections for similar analyses (e.g. (Liu et al., 2020)), and was deemed a reasonable selection by an expert.

We used brain-wide gene-expression data provided by the Allen Human Brain Atlas (AHBA). The AHBA dataset consists of 3702 tissue samples and respective microarray expression data from six human donors (Hawrylycz et al., 2012). For this dataset, the transcriptomic expression levels of more than 20,000 genes were measured in more than 50,000 probes across different cerebral and cerebellar regions. We accessed the AHBA database via the brainstat and abagen toolboxes (*Abagen: A Toolbox for the Allen Brain Atlas Genetics Data | Zenodo*, n.d.), which allows retrieving the transcriptomic data in Schaefer 400 parcellations. Brainstat fetches the tissue samples of all donors in MNI space and then applies intensity-based filtering, so that probes where more than 50% of samples exceeded a background noise threshold were excluded. It furthermore identifies differential stability across donors for probes indexing the same gene, and selects the most stable one. The remaining n = 15.631 genes were then matched to the respective Schaefer400 parcels and expression values were normalized across samples and genes with a scaled robust sigmoid normalization function. In a last step, expression values were averaged within each parcel, and then averaged again across all six donors. For our analysis, we only considered the left hemisphere of transcriptomic maps, where expression profiles for nearly all Schaefer parcels were available, whereas the right hemisphere lacks sufficient sample density as it had only been sampled for two out of the six donors.

We computed spearman correlations between gene expression enrichment for each of the selected GOIs with the observed differences in cortical microstructure between males and females using spin permutation tests. To avoid false-positive bias in spatial specificity tests of gene enrichment analysis due to spatial non-independence of brain maps, we tested for significant spatial overlap between the respective transcriptomic map relative to randomly spun phenotype maps (i.e. our effect maps of sex differences (Fulcher et al., 2021). For that, we adjusted the spin-test function from the ENIGMA toolbox, so that spherical representations of the unthresholded three phenotypic maps were randomly spun in 1000 permutations and correlated with the 25 transcriptomic maps of our GOIs (Alexander-Bloch et al., 2018) . This procedure accounts for spatial autocorrelations by leveraging the spherical representations of the cerebral cortex. We report the frequency in which the correlation between permuted phenotypic maps and genes exceeded the original test statistic as *spin-p-value*, as such correcting for family-wise correction for multiple comparisons. Lastly, to provide a measure of genetic specificity, we generated a measure of “brain-gene-baseline” and tested our effects against the baseline. We built the baseline transcriptomic map by extracting the principal component of all available transcriptomic maps in the left hemisphere. We provide spatial correlations (spearman) between phenotypic maps of sex differences in profile mean, skewness and gradients with the brain gene baseline as a reference.

### Histological decoding

To contextualize our findings histologically, we chose a theory-driven, manual histological characterization of “cortical types”, defined by con Economo (von Economo & Koskinas, 1925), digitalised to FSaverage space by Scholtens (Scholtens et al., 2018), and recently re-analysed by Garcia-Cabezas (García-Cabezas et al., 2020). Here, cortical areas are characterized according to different laminar features that are extracted from Nissl stained sections. Amongst others, the authors characterized the sharpness of boundaries between layers, prominence of deep (layers V and VI) or superficial (layers II and III) layers, degree of granularity of cells and presence of layer IV. Based on these features, the cortex was then divided into cortical types. The hierarchy from agranular cortex, dysgranular cortex, eulaminate cortex 1-3 and koniocortical areas is analogous to a hierarchy of more diffuse to more elaborate laminar differentiation of cortical histology. We assigned each Schaefer400 parcel to one of the six cortical types and ran the previously described spin-test based spearman correlations between cortical types and the identified sex differences for our three cortical microstructure measures.

### Control analyses: Vascular-hormonal coupling

Sex hormones influence structural plasticity as well as they modulate blood flow, vasodilation (Pelligrino & Galea, 2001), and haemoglobin concentration (Murphy, 2014). We had previously included ICV as a covariate in our models to account for potential hormone-related volume shifts in the brain, which may influence MR signal intensity (Lorio et al., 2016). However, since the T1w/T2w measure reflects water and iron content in the brain (Lorio et al., 2016; Stüber et al., 2014) as well as it does myeloarchitectural features (Glasser & Van Essen, 2011), we performed an additional control analysis post-hoc to our main analysis.

To identify which tissue changes drove the T1w/T2w hormonal effect, we investigated the spatial coupling of our effect maps with cortical vasculature maps. Veins and arteries in the brain are not distributed homogeneously, so that areas are differentially impacted by vasculature. We hypothesized that if the observed effects in T1w/T2w measures were mainly due to changes in hormone-related blood-flow changes, then areas in which the MRI signal is more strongly influenced by the vasculature should express the strongest effects. We used surface projections of an atlas which maps the distribution of arteries and veins in the brain (Bernier et al., 2018). To identify the influence of arteries and veins per parcel on the three microstructural measures, we first computed spatial spearman correlations between these two atlases and our main sex-difference effect maps. In a second step, we aimed at identifying the relationship between vasculature and the effect of hormones. We thus computed spatial correlations between hormonal-sub-group effect maps, and the two atlases, respectively. Again, to address the problem of statistical auto-correlation, we used our adjusted spin-test function for all spatial correlations, and built a zero-distribution out of spatial correlations between 1,000 randomly spun cortical spheres of the unthresholded t-statistics for sex differences for profile mean, profile skewness and the two cerebrovascular atlases, respectively.

## Author Contributions

S.K. and S.V. analyzed the data and created figures. S.B. pre-processed hippocampal data. S.K. wrote the manuscript. S.B., R.G.Z., A.S., B.B., S.W., H.L.S, J.S., S.E., and S.V. revised and finalized the manuscript for publication.

## Data availability

The data that support the findings of this study are available from the corresponding author upon reasonable request.

## Code availability

All code is available here.

## Footnotes

1 In this manuscript, the terms ‘female and male sex’ refers to self-reported sex. The authors appreciate the complexity of biological sex and the influence of gender on biology, and do not postulate a sex binary.

2 We classified individuals of female sex if they self-reported their gender as female and indicated that they are or have been menstruating in their lives.

