## Supplementary figures and tables for "Relating sex differences in cortical and hippocampal microstructure to sex hormones"

### Supplement

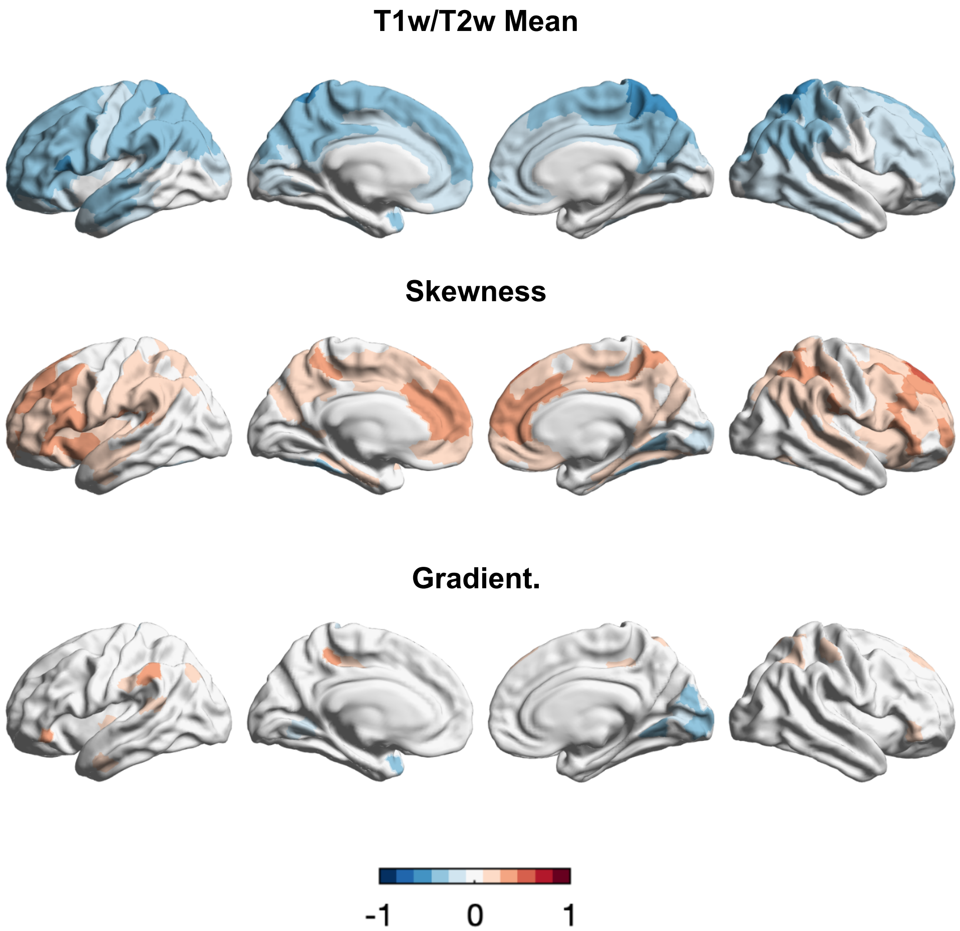


**Supplement 1. Intracortical T1w/T2w signal intensity profiling Additional control analyzes sex differences**. Shown are z-values for the female > males contrast, ontrolling for family structure (including the interaction between twin status and family status) and cortical thickness, FDR controlled Cohen’s d

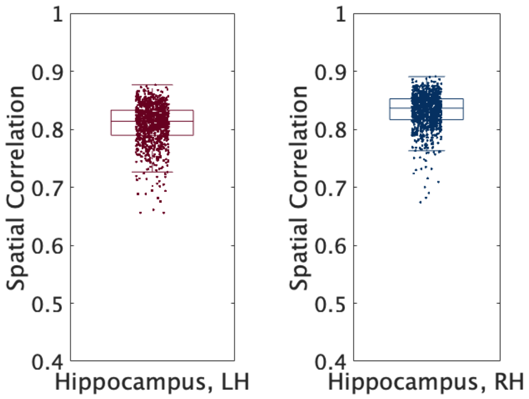

**Supplement 2. Internal Consistency Hippocampus**. Boxplots represent Pearson’s r-values between unthresholded t-statistics resulting from two respective split-halves of the sample (n = 1000 permutations) comparing the microstructural mean of the left and right hippocampus between females and males, indicating their reliability.

Low Estrogen

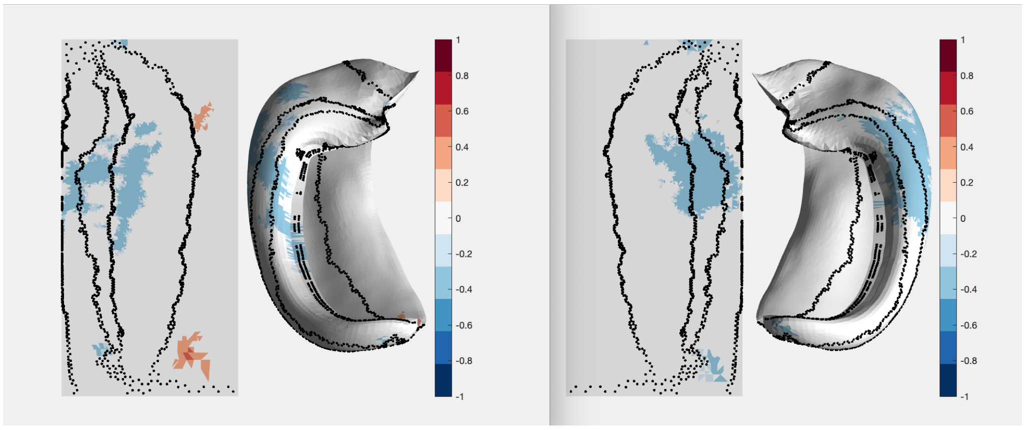


High Estrogen

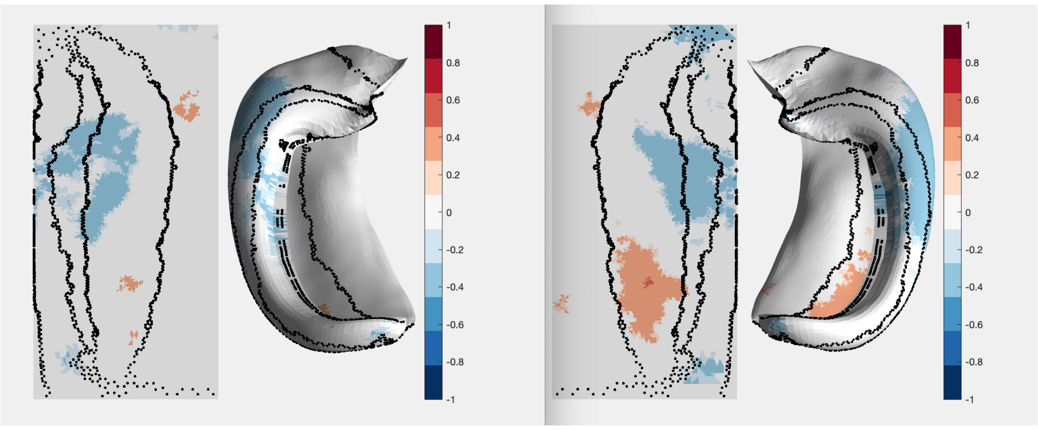


Low Progesterone

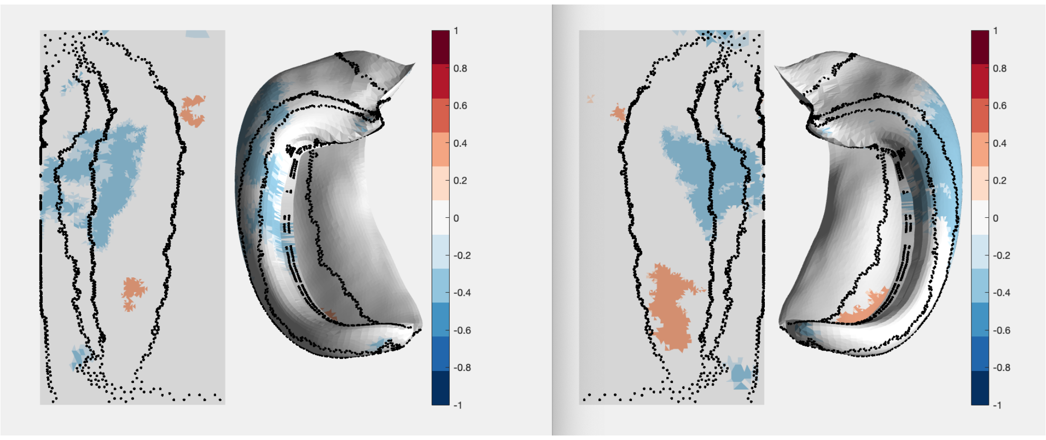

High Progesterone

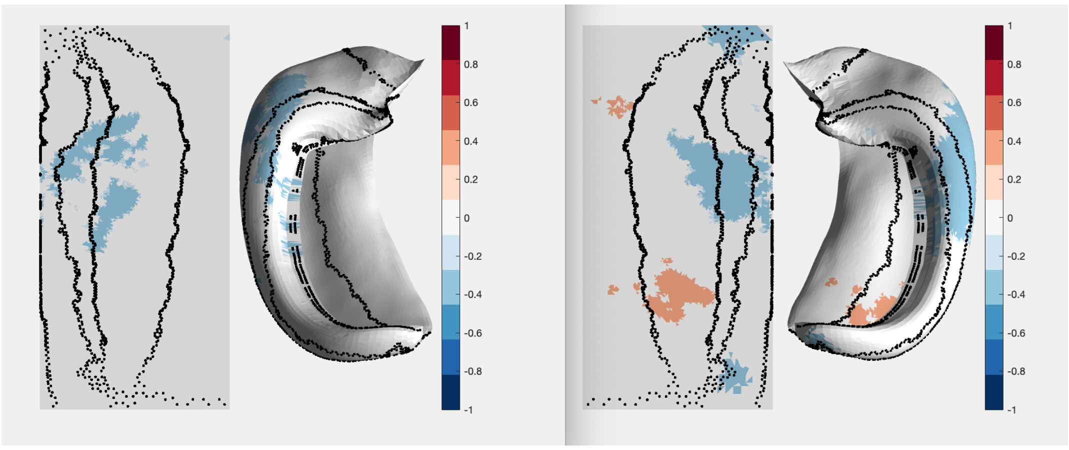


OC females

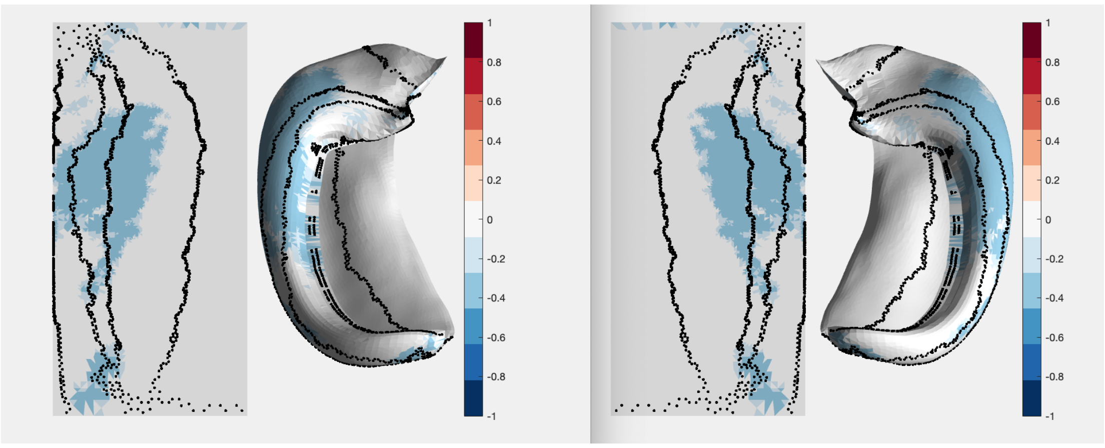


**Supplement 3. Hormonal effects in the hippocampus**. FDR-thresholded Cohen's *d* maps of T1w/T2w profile *mean* between males and subsamples of females divided by OC use and menstrual cycle phase projected on the cortical surface and the hippocampus.

**T1w/T2w mean**

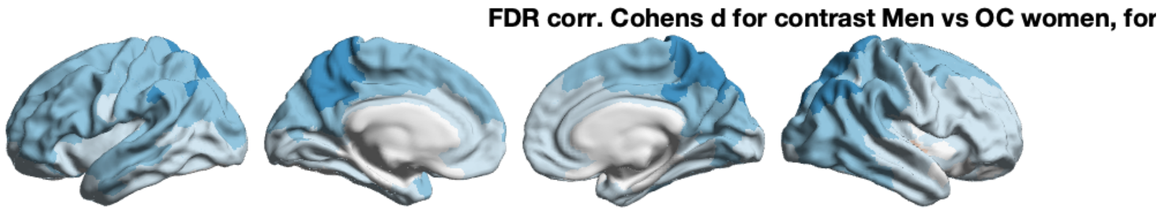

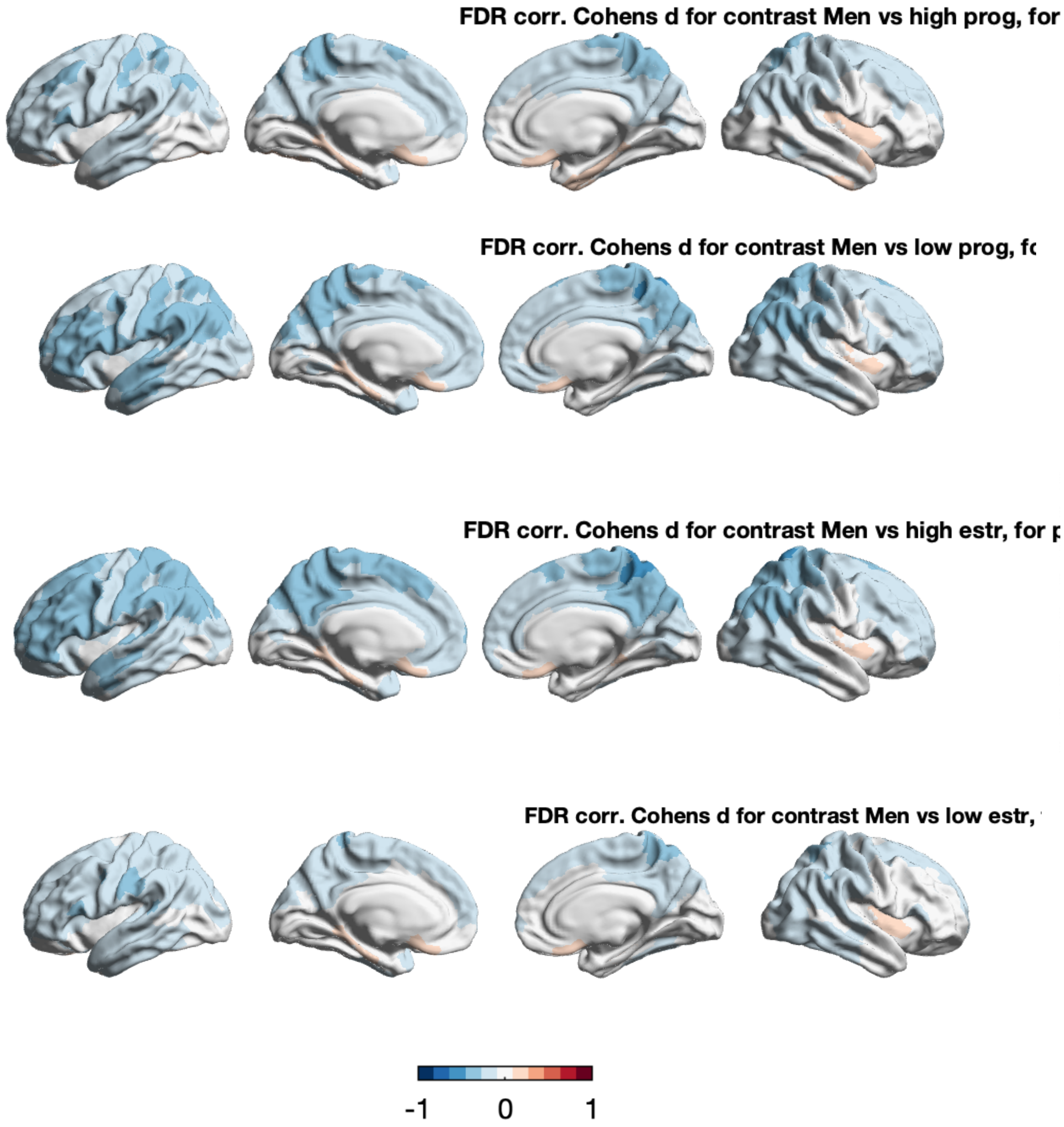


**T1w/T2w Skewness**

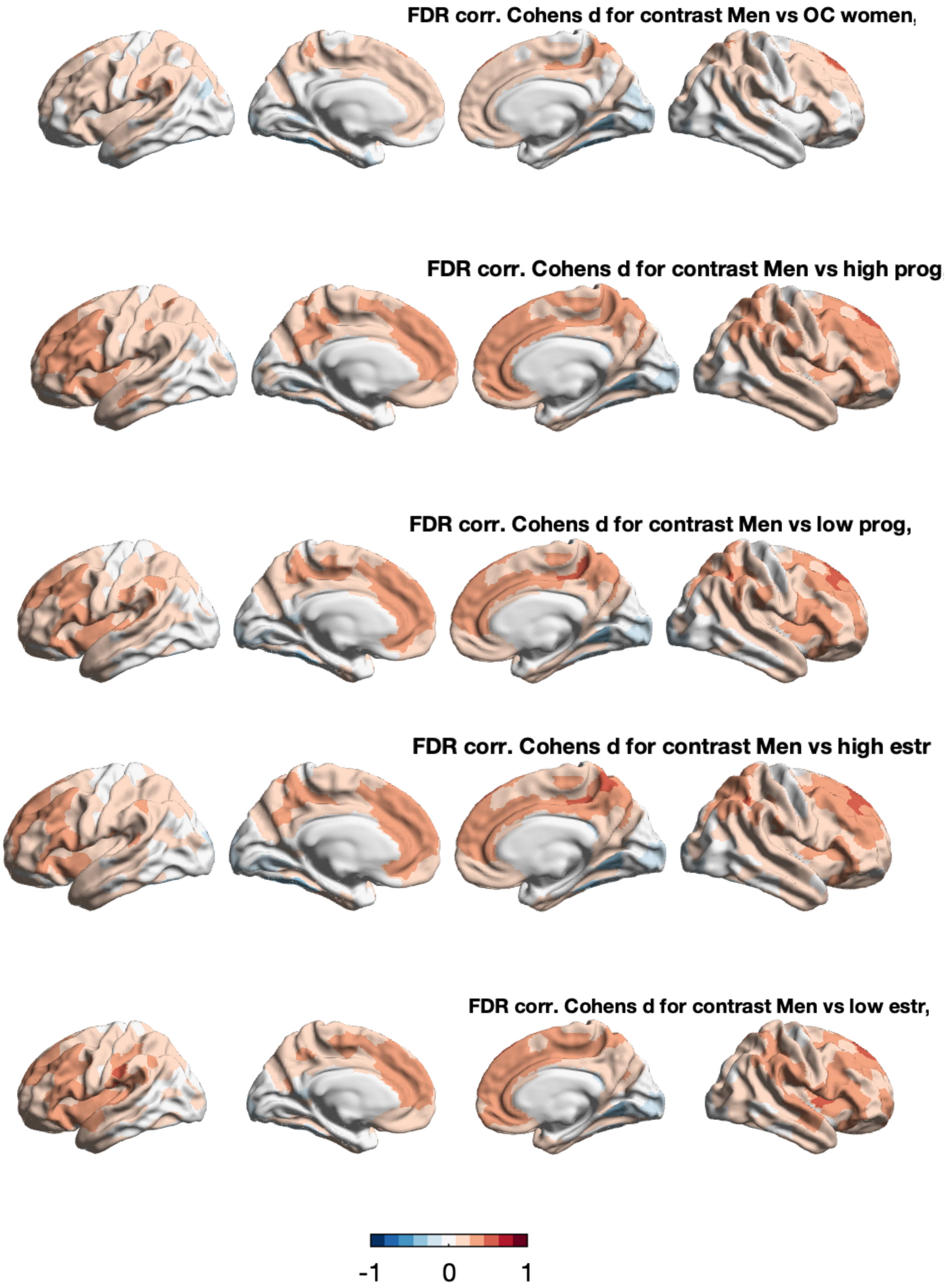
FDR corr. Cohens d for contrast Men vs OC women,
FDR corr. Cohens d for contrast Men vs high prog
FDR corr. Cohens d for contrast Men vs low prog,
FDR corr. Cohens d for contrast Men vs high estr
FDR corr. Cohens d for contrast Men vs low estr,
-1
0
1

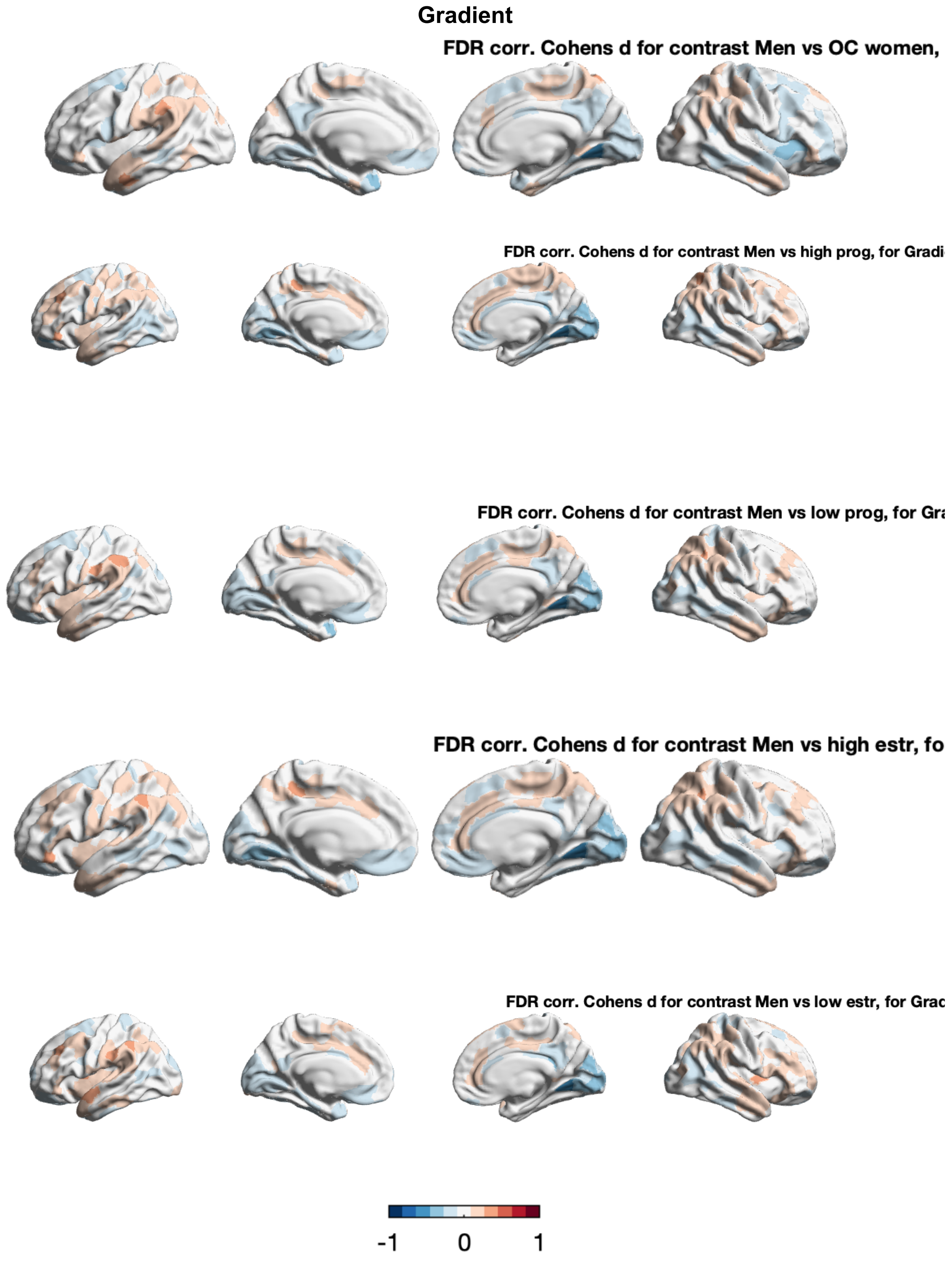


**Supplement 4. Hormonal effects in intracortical microstructure**. FDR-thresholded Cohen's *d* maps of T1w/T2w profile *mean* between males and subsamples of females divided by OC use and menstrual cycle phase projected on the cortical surface and the hippocampus. Shown is the females > males contrast for T1w/T2w mean, skewness and the gradient. The order is OC, high progesterone, low progesterone, high estrogen and low estrogen.

| Gene | Grad Corr | Grad P | Grad p spin | mean corr | mean p | mean p spin | skew corr | skew p | skew p spin |
|---|---|---|---|---|---|---|---|---|---|
| AR | -0.04 | 0.566 | 0 | -0.31 | 0.000 | 0.154 | -0.11 | 0.131 | 0 |
| ESR1 | -0.02 | 0.784 | 0 | 0.05 | 0.534 | 0 | -0.18 | 0.013 | 0.104 |
| ESRRA | -0.13 | 0.083 | 0 | -0.15 | 0.038 | 0.294 | -0.24 | 0.001 | 0.273 |
| ESRRB | -0.15 | 0.034 | 0.072 | 0.01 | 0.901 | 0 | -0.22 | 0.003 | 0.064 |
| ESRRG | -0.05 | 0.536 | 0 | -0.15 | 0.034 | 0.371 | -0.22 | 0.002 | 0.266 |
| GREB1 | -0.02 | 0.738 | 0 | -0.08 | 0.283 | 0 | -0.24 | 0.001 | 0.029 |
| PAQR6 | -0.01 | 0.918 | 0 | -0.05 | 0.501 | 0 | -0.02 | 0.835 | 0 |
| PAQR7 | -0.06 | 0.410 | 0 | 0.00 | 0.947 | 0 | -0.07 | 0.366 | 0 |
| PAQR8 | -0.16 | 0.028 | 0.121 | -0.17 | 0.021 | 0.128 | -0.03 | 0.730 | 0 |
| PAQR9 | -0.15 | 0.043 | 0.113 | 0.12 | 0.102 | 0 | -0.20 | 0.007 | 0.139 |
| PGRMC 1 | 0.11 | 0.135 | 0 | 0.26 | 0.000 | 0.196 | 0.18 | 0.015 | 0.232 |
| PGRMC 2 | -0.03 | 0.692 | 0 | 0.14 | 0.064 | 0 | 0.03 | 0.703 | 0 |
| AKR1C3 | 0.12 | 0.110 | 0 | -0.29 | 0.000 | 0.103 | 0.31 | 0.000 | 0.09 |
| CYP11A 1 | 0.08 | 0.279 | 0 | -0.04 | 0.601 | 0 | 0.11 | 0.117 | 0 |
| CYP17A 1 | -0.08 | 0.262 | 0 | 0.04 | 0.587 | 0 | -0.09 | 0.202 | 0 |
| HSD17B 1 | -0.11 | 0.142 | 0 | 0.08 | 0.249 | 0 | 0.02 | 0.749 | 0 |
| HSD17B 12 | -0.02 | 0.809 | 0 | -0.11 | 0.147 | 0 | 0.01 | 0.860 | 0 |
| HSD17B 3 | -0.12 | 0.101 | 0 | 0.13 | 0.077 | 0 | 0.01 | 0.863 | 0 |
| HSD17B 6 | 0.13 | 0.070 | 0 | -0.17 | 0.023 | 0.138 | 0.21 | 0.004 | 0.116 |

| | | | | | | | | | |
|---|---|---|---|---|---|---|---|---|---|
| HSD17B 7 | -0.12 | 0.087 | 0 | 0.12 | 0.112 | 0 | 0.00 | 0.989 | 0 |
| HSD17B 8 | 0.00 | 0.986 | 0 | -0.03 | 0.644 | 0 | 0.17 | 0.019 | 0.19 |
| PIBF1 | -0.03 | 0.641 | 0 | 0.00 | 0.999 | 0 | -0.25 | 0.000 | 0.038 |
| SRD5A1 | -0.14 | 0.062 | 0 | 0.09 | 0.215 | 0 | -0.15 | 0.042 | 0.166 |
| SRD5A3 | 0.01 | 0.910 | 0 | 0.31 | 0.000 | 0.064 | -0.01 | 0.872 | 0 |
| STS | -0.03 | 0.679 | 0 | -0.14 | 0.056 | 0 | -0.20 | 0.006 | 0.182 |

**Supplement 5. Genetic decoding, all results**. Spatial overlap between effect maps of sex differences for the microstructural gradient, profile mean and profile skewness**.** The table shows correlations, p values, and spin-corrected p-values. Transcriptomic maps of genes are of the following categories: sex hormone synthesis related genes, androgen receptor related, estrogen receptor related genes, and progesterone receptor related genes.
